# Nanopore sequencing unveils the complexity of the cold-activated murine brown adipose tissue transcriptome

**DOI:** 10.1101/2022.12.14.520420

**Authors:** Christoph Andreas Engelhard, Sajjad Khani, Sophia Derdak, Martin Bilban, Jan-Wilhelm Kornfeld

## Abstract

Alternative transcription increases transcriptome complexity by expression of multiple transcripts per gene and thus fine tunes cellular identity and function. Annotation and quantification of transcripts at complex loci using short-read sequencing is non-trivial. Recent long-read sequencing methods such as those from Oxford Nanopore Technologies (ONT) and Pacific Biosciences aim at overcoming these problems by sequencing full length transcripts. Activation of BAT thermogenesis involves major transcriptomic remodelling and positively affects metabolism via increased energy expenditure and endocrine factors. Here we comprehensively benchmark features of ONT long-read sequencing protocols compared to Illumina shortread sequencing assessing alignment characteristics, gene and transcript detection and quantification, differential gene and transcript expression, transcriptome reannotation and differential transcript usage (DTU). We find that ONT sequencing is superior to Illumina for transcriptome reassembly and reduces the risk of false-positive events due to the ability to unambiguously map reads to transcripts, at the expense of statistical power for calling differentially expressed features. We identified novel isoforms of genes undergoing DTU in cold-activated BAT including Cars2, Adtrp, Acsl5, Scp2, Aldoa and Pde4d, validated by RT-qPCR. Finally, we provide a reannotation of the murine iBAT transcriptome as a valuable resource for researchers interested in the molecular biology underlying the regulation of BAT.

## 4. Introduction

Alternative transcription (AT) is a post-transcriptional process in which multiple transcripts arise from a single gene locus by using alternative transcription start sites, altered polyadenylation sites and alternative splicing, thereby increasing the transcriptomic and translatomic complexity in a cell (de Klerk & ‘t Hoen, 2015). Alternative transcription is estimated to occur within 92% to 94% of human genes, substantially expanding the catalogue of co-expressed mRNAs (Pan et al., 2008; E. T. Wang et al., 2008). In line, sequencing of ribosome attached translated mRNAs (Ribo-seq) and proteomics studies confirmed that many RNA species produced by AT are translated and contribute to increased proteome diversity (Floor & Doudna, 2016; Lau et al., 2019; Rodriguez et al., 2020; Weatheritt et al., 2016). Interestingly, AT is tissue specific (Rodriguez et al., 2020; Xu et al., 2002) or marks specific cellular states (Fiszbein & Kornblihtt, 2017; Robinson et al., 2021), indicating a seminal role for AT in regulation of cellular identity and function.

Adipose tissue depots can be broadly classified into brown and white depots: While white adipocytes mainly function to store energy as triglycerides in large unilocular lipid droplets and coordinate energy metabolism by secretion of endocrine factors, brown adipocytes are densely packed with mitochondria and morphologically present multiple small lipid droplets (Rosen & Spiegelman, 2014). Upon sympathetic nervous system activation e.g., upon cold stimulus, brown adipocytes upregulate lipolysis, where the ensuring free fatty acids activate Uncoupling Protein 1 (UCP1) and generate heat by increasing the uncoupling of oxidative respiration from ATP generation. Additionally, sympathetic activation of brown adipocytes (BA) residing in brown fat and in inguinal white adipose tissue (so-called brown-in-white or ‘brite’ adipocytes) leads to profound changes in gene expression (Cannon & Nedergaard, 2004). Recent evidence suggests, that not only differential gene expression, but also the regulation of differential transcript usage (DTU) by RNA-binding proteins i.e., changes in the relative abundance of transcripts originating from one gene, is crucial for e.g., the regulation of adipocyte thermogenesis (Vernia et al., 2016). Transcriptomic studies have shown that DTU is required for adipogenesis, the process of differentiation of preadipocytes into mature adipocytes (Fiszbein & Kornblihtt, 2017; Yi et al., 2020). Moreover, AT events in key brown adipocyte genes such as the transcription factors *Pparg* and *Prdm16* have been reported to play a role in the control of brown adipocyte function (Chi & Lin, 2018; D. Li et al., 2016).

However, most studies focussing on AT and DTU so far have used Illumina short-read sequencing. Short-read sequencing inherently underperforms in relation to assembling transcripts, since the single reads only span a fraction of a transcript, and therefore requires complex computational post processing for transcriptome reassembly. This poses a conceptual problem: If two individual AT events in one gene occur too far away from each other to be spanned by a single short read, it is challenging to unambiguously decide if both AT events happen (i) in conjunction, (ii) arise independently from each (iii) or are mutually exclusive (Byrne et al., 2019). Short-read sequencing also suffers in respect to transcript level quantification required for analysis of DTU, as only reads mapping to parts of a gene unique to a single transcript can be unambiguously assigned to a transcript, while all others must be assigned based on statistical models (Bray et al., 2016; Patro et al., 2017). Long-read sequencing methods such as those developed by Pacific Biosciences (Eid et al., 2009; Sharon et al., 2013) and Oxford Nanopore Technologies (ONT; Garalde et al., 2018) generate full-length isoform reads that mitigate these limitations, allowing for simple transcriptome reannotation and unambiguous read assignment (Stark et al., 2019). Importantly, thousands of novel transcripts across a large collection of different human tissues have recently been revealed using long-read sequencing with ONT, enabling an understanding of functionally distinct protein isoforms that different transcripts can give rise to (Glinos et al., 2022). Reference databases like GENCODE are based on a limited number of tissue transcriptomes (Robinson et al., 2021; Xu et al., 2002). Since alternative transcription is tissue and cell state specific, it is of high biological interest to reannotate transcriptomes in cell types such as brown adipocytes, which are not represented in reference annotations, in order to identify and quantify tissue specific transcript isoforms. Long-read sequencing methods like ONT sequencing on the other hand, suffer from lower throughput and lower base calling accuracy, resulting in failure to detect lowly expressed isoforms and fuzzy splice junction annotation (Stark et al., 2019). Accordingly, algorithms that combine short and long reads for improved transcriptome reassembly, either *de novo* or using a reference genome, have been developed (Fu et al., 2018; Kovaka et al., 2019; Tang et al., 2020) and consortia such as GENCODE have started incorporating long-read sequencing in their reference transcriptome annotation pipelines (Frankish et al., 2021).

Here, we have compared three different library preparation methods using the ONT platform and assessed their ability for transcript detection, quantitation and differential expression calling in addition to performing transcriptome reassembly and analysis of differential transcript usage. Using RNA isolated from murine iBAT, we identified cold induced isoform switches in genes regulating thermogenic β3 adrenergic receptor (AR) signalling at multiple levels including regulation of cAMP levels (*Pde4d*) and receptor signalling (*Adtrp*), lipid metabolism/signalling (*Scp2, Mlixpl*) and protein sorting (*Ergic1*). Finally, using FLAIR/ChIP Seq we identified a novel alternative transcription start site in the in the mitochondrial respiration regulating protein *cysteinyl-tRNA synthetase gene (Cars2)* and validated coding potential for an alternative (shorter) transcript (*Cars2*-AT) using the *Coding-Potential Assessment* (CPA) Tool and determined functional domain structure using pfam. As an example which demonstrates the potential of the ONT long read iBAT transcriptome reannotation reported here, we show that sgRNAs targeting the Cars2-AT promoter, are efficient in inducing the expression of Cars2-AT in brown adipocytes *in vitro*. Thus, we provide a reannotation of the murine iBAT transcriptome which can be a valuable resource for researchers interested in iBAT biology by facilitating them to target the relevant isoforms of a gene in study and detect novel DTU events in cold activated murine iBAT, demonstrating the contribution of AT in the regulation of brown adipocyte activity.

## 5. Results

### Comparison of Nanopore-based approaches for transcript resolution of cold-activated BAT

We isolated RNA from iBAT of 20-week old, male C57BL/6N mice cold treated for 24 h at 4 °C or housed at room temperature (n = 3). To evaluate different library preparation methods for the ONT sequencing platform, we prepared libraries using *(i)* direct cDNA sequencing, which is PCR-free and avoids bias introduced by the amplification (Chen et al., 2021; Stark et al., 2019) and *(ii)* TeloPrime sequencing, which uses a 5’ cap specific template switching oligo to enrich for full-length RNA but requires PCR amplification. All samples were multiplexed, and library pools sequenced on two separate flow cells per library preparation method on a ONT GridION to assess the variability in performance of the flow cells. Additionally, the samples were pooled within the respective treatment group and *(iii)* sequenced following ONT’s direct RNA protocol on one flow cell each. Finally, *(iv)* we performed strand specific, paired-end short-read sequencing following Illumina’s TruSeq protocol as reference (Fig. 1A). To the best of our knowledge, this analysis represents the most comprehensive characterization of full-length transcripts and transcript diversity to date in the murine BAT depot, both at basal levels and upon cold activation. Low quality ONT reads with a minimum average Phred Quality Score below 7 (20 % base call accuracy) were removed, leaving 13.5×10^6^ reads from TeloPrime sequencing, 12,7×10^6^ reads from direct cDNA sequencing, and 3,0×10^6^ from the direct RNA sequencinq (Tab. 1). Interestingly, we observed a high variability in the number of high-quality reads produced per flowcell, varying by 41 % for direct cDNA-Seq and 16 % for TeloPrime-Seq, even though the same libraries were used, and flow cells run in parallel (Fig. 1B). Similar variability in MinION flow cell performance has been noticed in other studies (Ip et al., 2015; Oikonomopoulos et al., 2016; Sessegolo et al., 2019). Noteworthy, a striking difference in high quality read numbers was seen between the samples harvested from mice housed at room temperature compared to those from cold treated animals in TeloPrime-Seq. Read length distributions were similar between samples and flow cells within one library preparation method (Fig. 1C). In good agreement with other reports (Udaondo et al., 2021), average read lengths were similar for direct RNA-Seq (1,033 nt) and direct cDNA-Seq (1,141 nt) but longer for TeloPrime (1,326 nt). The read length distribution of the TeloPrime method however was multimodal, while the other distributions were unimodal. Average read quality was similar between samples, but interestingly depended on the flow cell used (Fig. 1D), in agreement with the number of high-quality reads received. Thus, our comparisons show that the TeloPrime protocol enriches for longer RNA molecules compared to other long-read protocols, but also reveal substantial technical variation between flow cells.

**Tab. 1.**
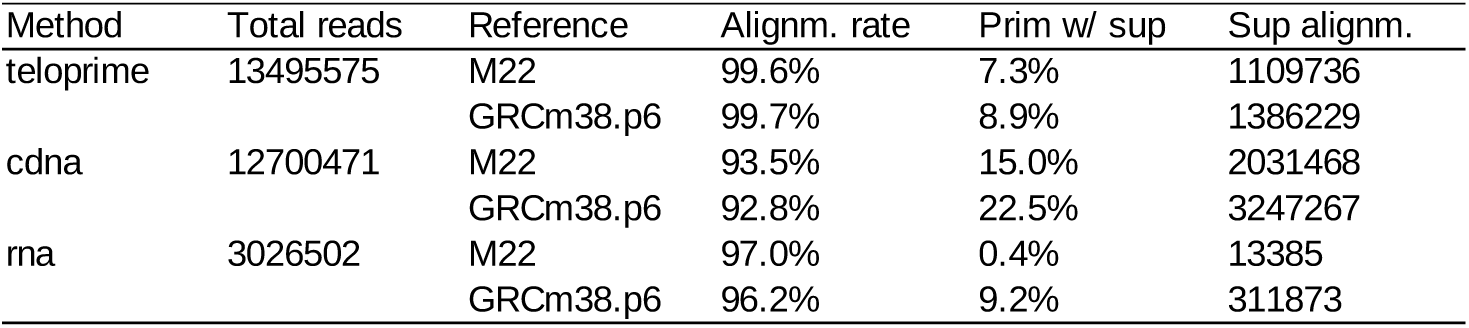
Alignment characteristics. Number of total reads (Q > 7), alignment rates, rate of reads with supplementary alignments and number of supplementary alignments for the three long-read sequencing methods, stratified by mapping to the genome (GRCm38.p6) or transcriptome (GENCODE M22).

| Method | Total reads | Reference | Alignm. rate | Prim w/ sup | Sup alignm. |
| --- | --- | --- | --- | --- | --- |
| teloprime | 13495575 | M22 | 99.6% | 7.3% | 1109736 |
|  |  | GRCm38.p6 | 99.7% | 8.9% | 1386229 |
| cdna | 12700471 | M22 | 93.5% | 15.0% | 2031468 |
|  |  | GRCm38.p6 | 92.8% | 22.5% | 3247267 |
| rna | 3026502 | M22 | 97.0% | 0.4% | 13385 |
|  |  | GRCm38.p6 | 96.2% | 9.2% | 311873 |

**Fig. 1.**
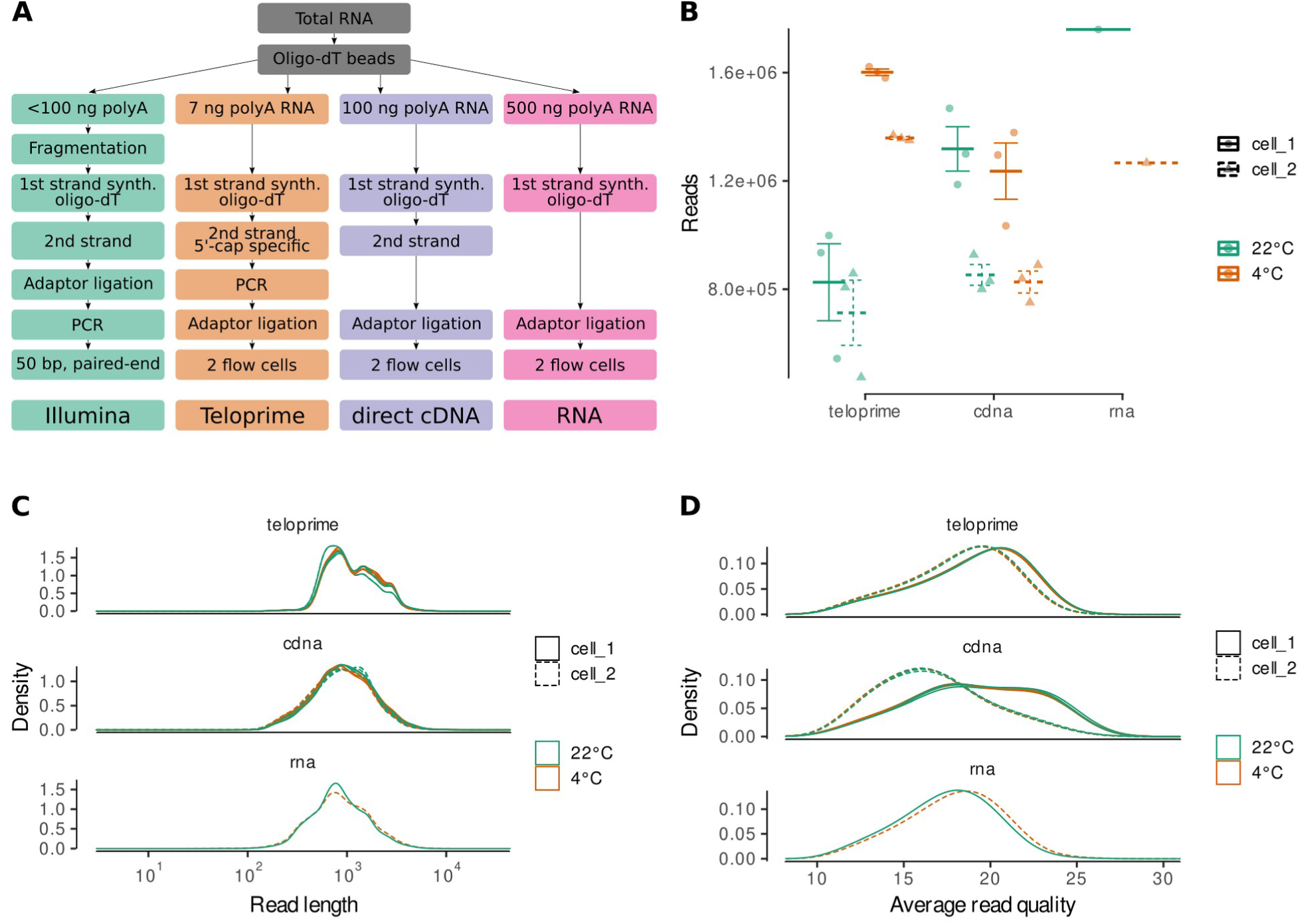
Characterisation of ONT reads. **A** Experimental design. RNA was either sequenced by the Illumina short-read platform or by three different ONT long-read library preparation protocols: TeloPrime, direct cDNA sequencing (cdna) or direct RNA sequencing (rna). See results and methods for details. **B;C;D** Total number of reads (B), read length distribution (C) and read quality distribution (D) by ONT sequencing method, flow cell and housing temperature.

### ONT teloprime improves coverage of full-length transcripts

We next aligned the quality filtered long reads to the murine genome and transcriptome using minimap2. Overall, alignment rates were high and independent of whether the alignment was performed using the transcriptome or genome as reference, ranging from 93 % for the direct cDNA-Seq method to > 99 % for TeloPrime (Tab. 1), emphasizing one of the main advantages for long-read RNA-seq. In line with other reports, we noted accompanying supplementary alignments i.e., reads not mapping linearly to the reference (Tab. 1; Fig. 2A; Fig. S1A-C). Reads with supplementary alignments were most common in direct cDNA-Seq and these were longer (Fig. 2C; Fig. S1E) and showed larger unaligned parts (Fig. 2D; Fig. S1F) then reads from the other methods. Supplementary alignments can arise from reads mapping to different chromosomes, indicative for chromosomal rearrangements (Klever et al., 2020; Leung et al., 2021). However, in direct cDNA-Seq, supplementary alignments mostly mapped anti-sense to the same transcript as the primary alignment (Fig. 2B; Fig. S1D), indicating that the second strand of the cDNA was sequenced subsequently to the first strand. Next, we compared the ability of the different long-read sequencing methods to cover full transcripts. Comparison of the read length distribution of the different methods to the hypothetical distribution of transcript lengths as inferred from Illumina-Seq transcript abundances revealed that the direct cDNA-Seq method generated a read length distribution shifted towards shorter reads (Fig. S1G). This was less prominent for the TeloPrime and direct RNA-Seq methods. Investigation of gene body coverages revealed that Illumina short-read sequencing mostly covers the middle part of transcripts, with reduced coverage at the 3′ and 5’ of the gene body in comparison to ONT sequencing, as demonstrated previously (Fig. 2E; Leshkowitz et al., 2022; Soneson et al., 2019; Wright et al., 2022). All long-read libraries showed decreasing coverage from the 3’ end of transcripts towards the 5’ end. Of note, this decrease was markedly reduced in TeloPrime-Seq, which is meant to enrich for full length transcripts. To assess the fraction of transcripts covered by reads and the proportion that represent full-length transcripts, a coverage fraction was calculated. We defined coverage fraction as the observed transcript length (alignment length) divided by the original known transcript length. We observed that both the coverage fraction as well as the fraction of full length reads markedly decreased with transcript length, in line with previous reports (Soneson et al. 2019; Fig. 2F; Fig. S1H). This could have been caused by RNA degradation during library preparation protocols or software artefacts during the base-calling process (Kovaka et al., 2019; Sessegolo et al., 2019). Overall, coverage was higher in TeloPrime-Seq as reported (Sessegolo et al., 2019). Direct cDNA-Seq showed both the lowest coverage and the smallest fraction of full length reads. As expected, the coverage of supplementary mappings was lower compared to primary alignments, although full-length supplementary alignments were present in all three ONT methods. Thus, we demonstrate that the TeloPrime protocol enriches for longer, potentially full length, transcripts compared to other two full-length sequencing protocols due to its dedicated library preparation method.

**Fig. 2.**
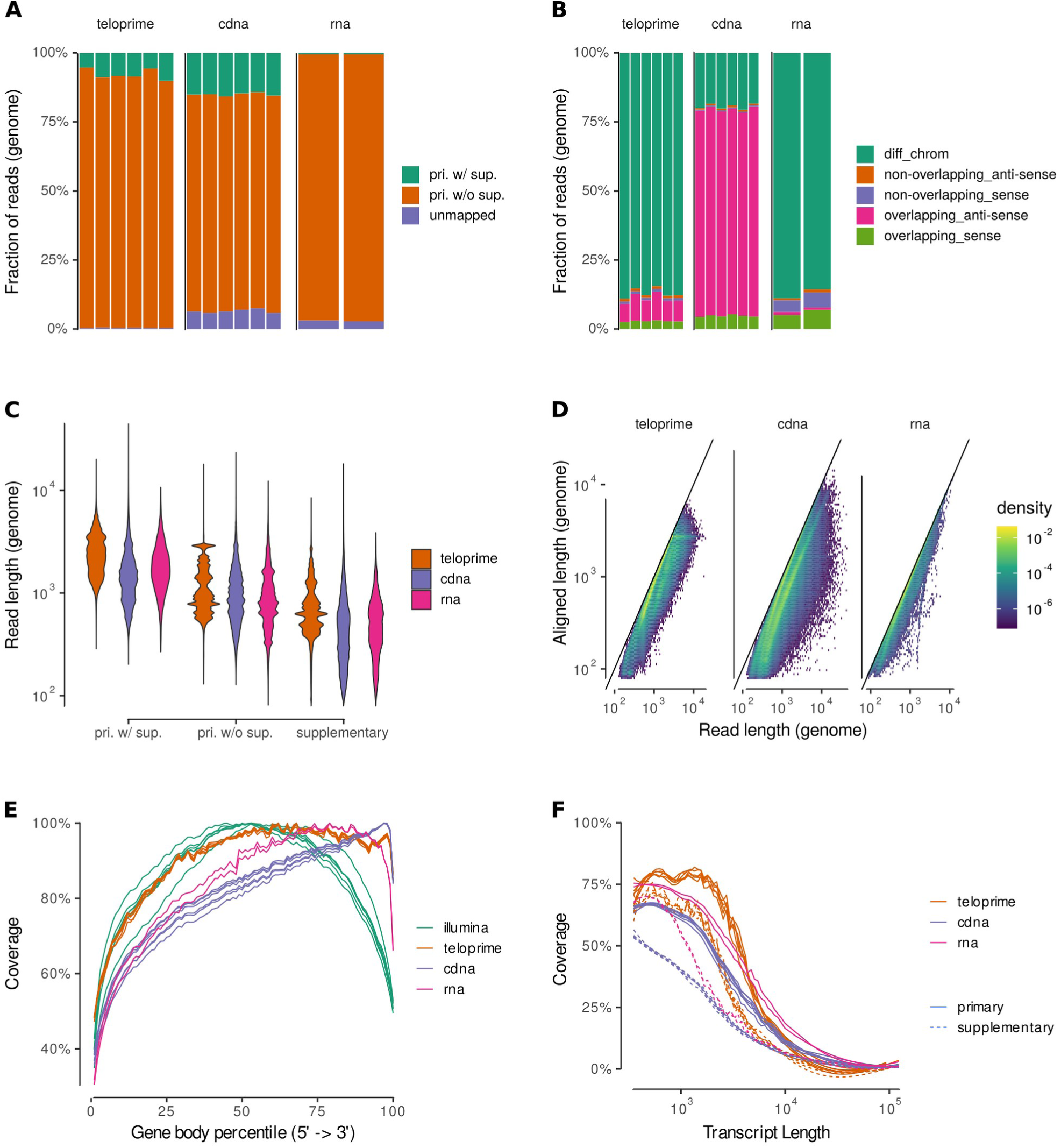
Characterisation of read alignments. **A** Fraction of reads classified by whether the primary alignment against the genome has at least one supplementary alignment attached to it. **B** Fraction of supplementary alignments against the genome stratified by their relation to the corresponding primary alignment. **C** Read length distribution for reads aligned to the genome. **D** Aligned length vs read length for primary alignments to the genome. **E** Percentile wise coverage of the gene body based on primary genome alignments. **F** Smoothed average transcript coverage of alignments mapped to the transcriptome for transcripts > 350 nt.

### ONT direct RNA and cDNA protocols show less bias for gene/transcript detection compared with TeloPrime

We next compared the ability of the different sequencing methods to detect expressed genes and transcripts based on the reference annotation. In agreement with previous reports (Sessegolo et al., 2019), direct cDNA and RNA-Seq detected a similar number of features as short-read sequencing at any given sequencing depth far outperforming TeloPrime-Seq (Fig. 3A and B). While differences in sequencing depth explain the large sets of genes detected by Illumina-Seq or by direct cDNA-Seq and Illumina-Seq alone, we also observed 1447 genes only in direct RNA-Seq, indicating technical biases of the different methods. Similarly, most transcripts were observed in direct cDNA and Illumina-Seq. However, the share of transcripts only detected using one but not the other protocol was even more pronounced then on gene level, indicating differences in the transcript identification potential of the different technologies (Fig. 3C). To detect the cause of these differences, we stratified the transcript detection rates by transcript length and transcript biotype (Fig. 3D). As reported, detection rates for Illumina, direct cDNA-Seq and direct RNA-Seq increased with transcript size (Soneson et al., 2019). TeloPrime-Seq detection rates on the other hand were highest for transcripts ranging from 1000 nt to 3000 nt. Detection rates of the direct cDNA-Seq method reached the detection rates of short-read sequencing for coding genes longer than 5000 nt but not for long noncoding RNAs (lncRNAs). Genes and transcripts detected by either long-or short-read sequencing alone were enriched for noncoding RNA as compared to those detected by both sequencing Fig. S2A,B). Since coding genes are generally higher expressed than other classes of RNA (Cabili et al., 2011), we assessed a potential effect of expression level on the gene and transcript detection rates by the ONT methods (Fig. 3E; Fig. S2C). Both genes and transcripts detected by short-read sequencing and long-read sequencing showed on average a higher expression measured by Illumina-Seq compared to features detected by short-read sequencing only. However, there were also highly expressed genes and transcripts that were not detected by the ONT sequencing protocols (Fig. S2D;E). Interestingly, there were also features with high expression in the ONT datasets that were not detected by short-read sequencing, more prominently for transcripts compared to genes. Thus, direct cDNA-Seq and direct RNA-Seq had comparable gene and transcript detection rates, which were proportional to gene and transcript length, whilst TeloPrime yielded lower detection rates with a non-linear relationship to transcript length.

**Fig. 3.**
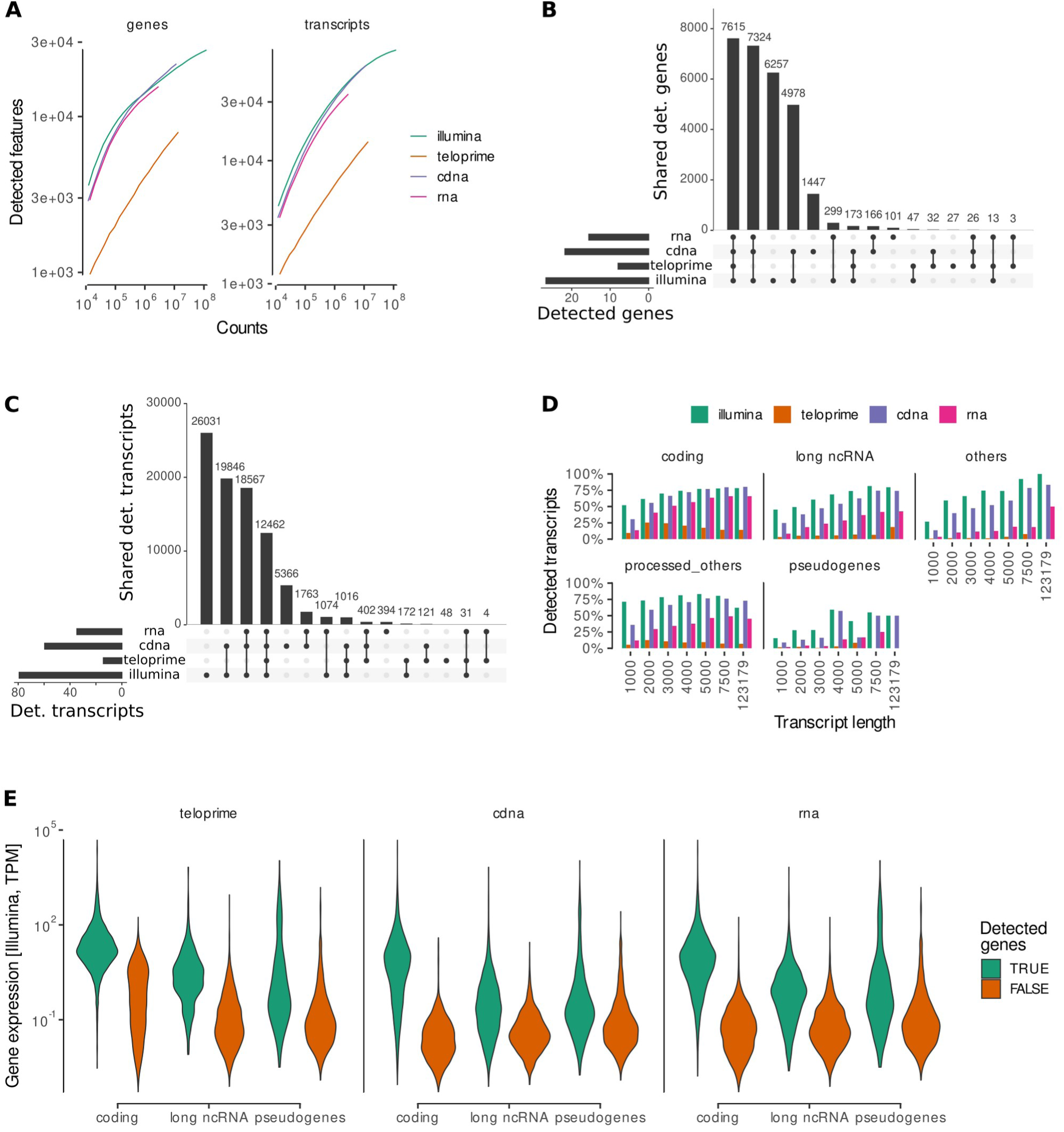
Feature detection. **A** Feature detection rate by library size. A feature is counted as detected if there is at least one primary alignment to it. **B;C** Overlap of detected genes (B) and transcripts (C) between the different sequencing methods. **D** Transcript detection rate by transcript length and biotype. 100 % is any transcript detected in any of the sequencing datasets. E Abundance in the Illumina dataset of genes either detected or not by the different ONT library preparation methods.

### Direct RNA- and cDNA-Seq is superior to TeloPrime for gene/transcript quantification

RNA-Seq and cDNA-Seq correlated very well on gene (*R^2^* = 0.92; Fig. 4) and transcript level (*R^2^* = 0.92; Fig. S4A). However, the situation was different when comparing the long-with short-read sequencing: While direct cDNA and RNA-Seq results correlated well with the abundance measured by Illumina sequencing on gene level (*R^2^* = 0.85 and 0.87), larger differences occurred on transcript level (*R^2^* = 0.54, both). The estimation of transcript abundance is challenging as transcripts from one gene share large parts of their sequence, causing ambiguity in read assignments when using short-reads (Soneson et al., 2016). TeloPrime-Seq quantification correlated less with the other methods. Noteworthy, the slope of the ratio of TeloPrime counts to those of other methods was larger than 1, indicating that the TeloPrime method overestimates the expression of highly abundant features and underestimates lowly expressed features, an observation also made in ONT RNA-Seq using PCR amplification (Chen et al., 2021). As we sequenced all TeloPrime and direct cDNA samples on two different flow cells, we could make use of technical replicates to assess the variability in sequencing performance between different flow cells. While sequencing counts for TeloPrime sequencing correlated very well among the two flow cells (*R^2^* = 0.86 to 0.90; Fig. S3B), the direct cDNA-Seq method showed a higher variation (*R^2^* = 0.75 to 0.77; Fig. S3C) reflecting the variability in read quality and length distributions (Fig. 1B-D). Thus, we find that direct RNA and direct cDNA protocols are most similar, reflecting a more unbiased representation of the transcriptome in comparison to TeloPrime.

**Fig. 4.**
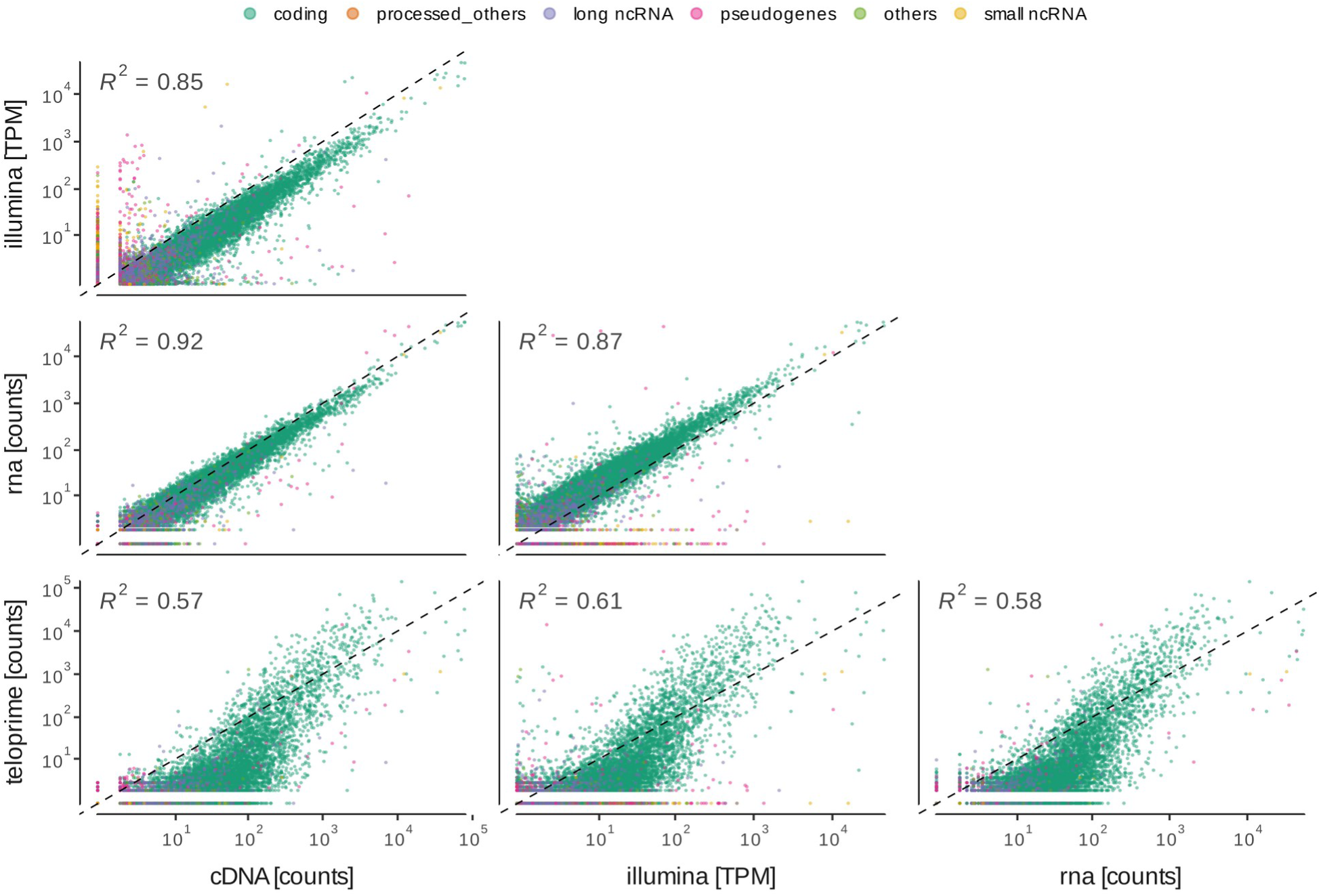
Gene quantification. Scatter plots showing the correlation in gene quantification between the different sequencing methods.

### Differential gene and transcript expression analysis in cold-activated BAT

The main goal of feature quantification is to detect differentially expressed genes and transcripts across biological samples. We compared the performance of the different ONT library preparation methods and Illumina sequencing to detect such features between iBAT of mice housed for 24 h either at room temperature or 4°C. Overall, the largest number of differentially expressed genes and transcripts (989 and 1195, respectively) was detected by Illumina sequencing followed by TeloPrime- (568 genes and 552 transcripts) and cDNA-Seq (489 genes and 476 transcripts) (Fig. 5A). Each sequencing method detected a unique set of features not seen by the other methods (312, 247 and 47 for Illimuna, TeloPrime and direct cDNA, respectively). Of note, irrespective of the method, most features identified as differentially expressed were protein coding genes. Interestingly, Illumina and direct cDNA performed better in detecting differentially regulated lncRNA genes compared with the TeloPrime protocol (Fig. 5B). Next, we compared the expression levels and fold changes between genes called to show differential gene or transcript expression by one of the ONT methods alone, Illumina sequencing alone or both methods. We found that genes differentially regulated according to Illumina but not the ONT methods showed low expression in both types of analysis, but more evident in the long-read method (Fig. 5C). On the other hand, genes detected to be differentially regulated by the ONT methods showed similar expression levels in both short- and long-read sequencing, independent of their status according to Illumina-Seq, but those features called by both methods showed higher fold changes in Illumina-Seq upon cold treatment. Thus, we observed that TeloPrime-Seq showed higher fold changes compared to Illumina-Seq, confirming that this method overestimates highly expressed and underestimates lowly expressed features, and that transcripts (but not genes) called by either of the long but not short-read sequencing methods showed lower expression on average in Illumina-Seq (Fig. S4).

**Fig. 5.**
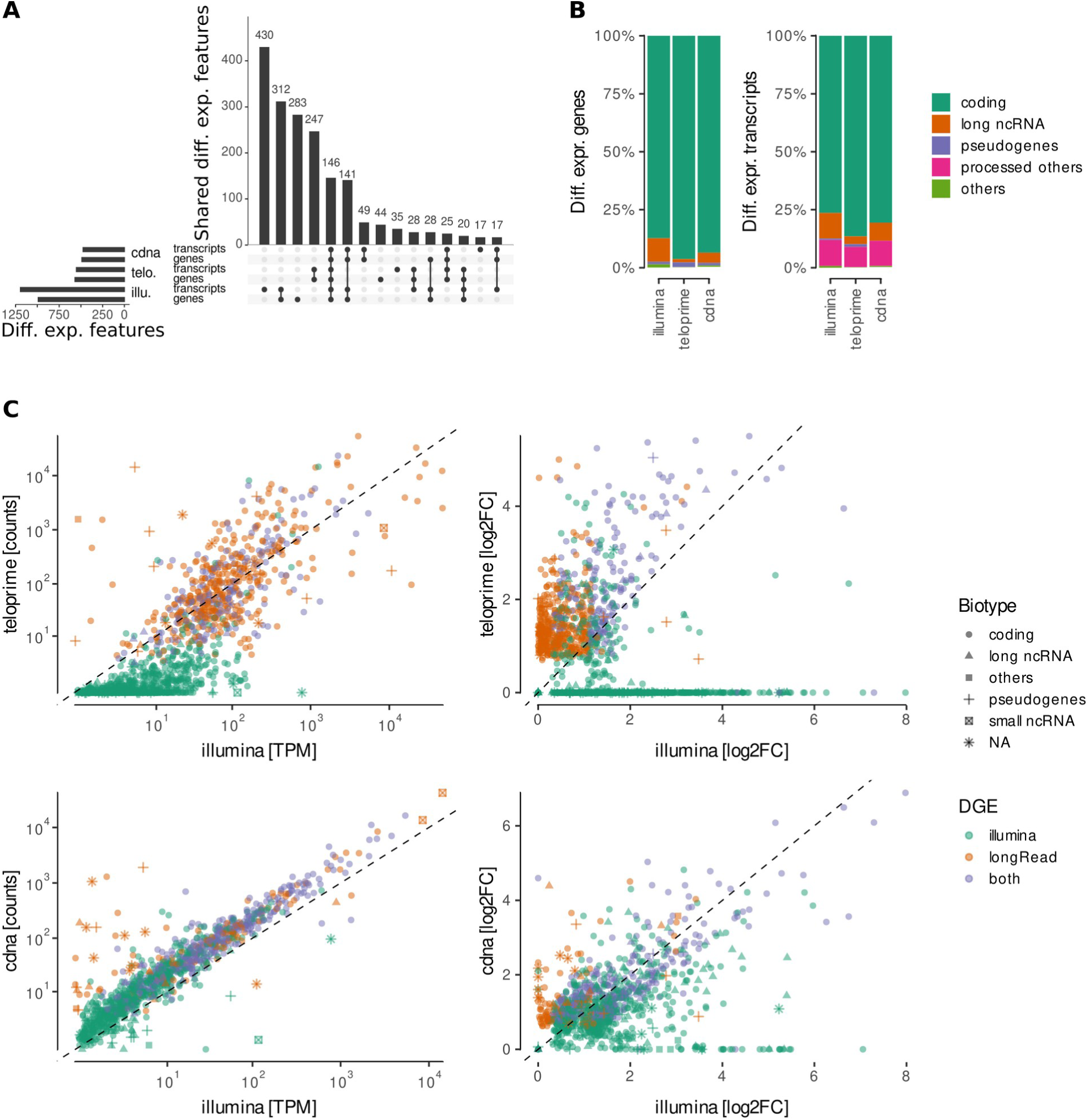
Differential gene expression analysis. **A** Overlap between genes showing differential gene expression or genes with at least one transcript showing differential transcript expression over the two ONT methods and Illumina sequencing. **B** Biotypes of genes and transcripts showing differential gene and transcript expression respectively compared between the different sequencing methods. **C** Comparison of expression levels and fold changes of genes showing significant differential expression between direct cDNA/TeloPrime and Illumina sequencing.

### ONT long read reannotation reveals novel features of the murine BAT transcriptome

The ability of long reads to unambiguously identify expressed isoforms facilitates the analysis of complex splicing events involving multiple exons. To reveal the nature and magnitude of newly identified transcripts in murine brown adipose tissue, we applied two transcriptome reassembly algorithms. FLAIR (Tang et al., 2020) corrects splice-sites of long-reads based on known, user-provided annotation e.g., from short reads, filters for those long-reads starting at given transcription start sites (TSS) and then collapses the long-reads to transcripts, keeping those with a minimum coverage of ONT reads. Stringtie (Pertea et al., 2015; Shumate et al., 2021) creates a splice graph based on long-reads, moves the splice junctions in this graph to the nearest junctions supported by short-read sequencing, removing them if not supported, and uses both short and long reads to filter for a minimum coverage. Each identified transcript was assigned to a structural category describing the type of relationship to the reference transcript (Fig. 6A). Generally, stringtie reannotated more transcripts compared to FLAIR (Fig. 6A), as shown previously (Kovaka et al., 2019). Irrespective of the reannotation algorithm, direct cDNA sequencing yielded the highest, while direct RNA sequencing gave the lowest number of reannotated transcripts, resembling the number of mapped reads (Fig. 3B-D), suggesting that transcript identification is affected by the ONT sequencing protocol. In all stringtie and the direct RNA-Seq FLAIR reannotation, most reannotated transcripts fully matched reference annotations (“full splice match”; Fig. 6A). Novel transcripts not present in the reference annotation (“novel in catalog”) made up for the second largest set and were relatively more prominent in FLAIR reannotations compared to stringtie reannotations, especially in the FLAIR-TeloPrime reannotation. Transcripts missing exons from either the 3’ or 5’ end (i.e. “incomplete splice match”; ISM) comprised the third largest class. Of note, the TeloPrime-(FLAIR) dataset was almost devoid of ISM transcripts, most likely because of its selective enrichment for full-length RNA molecules. Combinations of known splice junctions or splice sites were the prevailing mechanisms underlying transcript diversity among “novel transcripts” (Fig. 6B). Among the ISM, class 5’ fragments as well as mono-exonsmatches”, were most often found (Fig. 6C). Interestingly, ISMs were more prominent when the TeloPrime data were used for reannotation by stringtie, suggesting that the hybrid approach might reintroduce truncated isoform annotations potentially based on degraded RNA molecules. Noteworthy, even though only primary reads were used for the reannotation, the direct cDNA and the stringtie reannotation of the TeloPrime data, featured a substantial amount of antisense transcript annotations (Fig. 6A). These were on average shorter (Fig. S5A) and consisted of less exons compared to the full splice matching transcripts (Fig. S5B), indicating they might be artefacts. These annotations might interfere with transcript mapping, especially for non-directional sequencing methods. Comparison of the overlap of reannotated reference transcripts between the different datasets showed that the largest sets of transcripts were either detected by all combinations of reannotation algorithm and sequencing method, or only in all the stringtie datasets, highlighting the strong impact of the transcriptome reassembly method (Fig. 6D). Intriguingly, large sets of transcripts were only detected in the teloprime/stringtie or in the Illumina dataset and to a smaller extent in the cDNA/stringtie and RNA/stringtie datasets. Among the FLAIR reannotations, only direct cDNA-Seq showed transcripts specific for this method. Curiously, while the TeloPrime/FLAIR dataset included the smallest amount of reference transcripts, the combination of TeloPrime and stringtie reannotated the highest number of reference transcripts apart from the Illumina sequencing-based reannotation. Thus, the ONT sequencing method had a significant impact not only on the number but also on the nature of the structural category of novel transcripts identified.

**Fig. 6.**
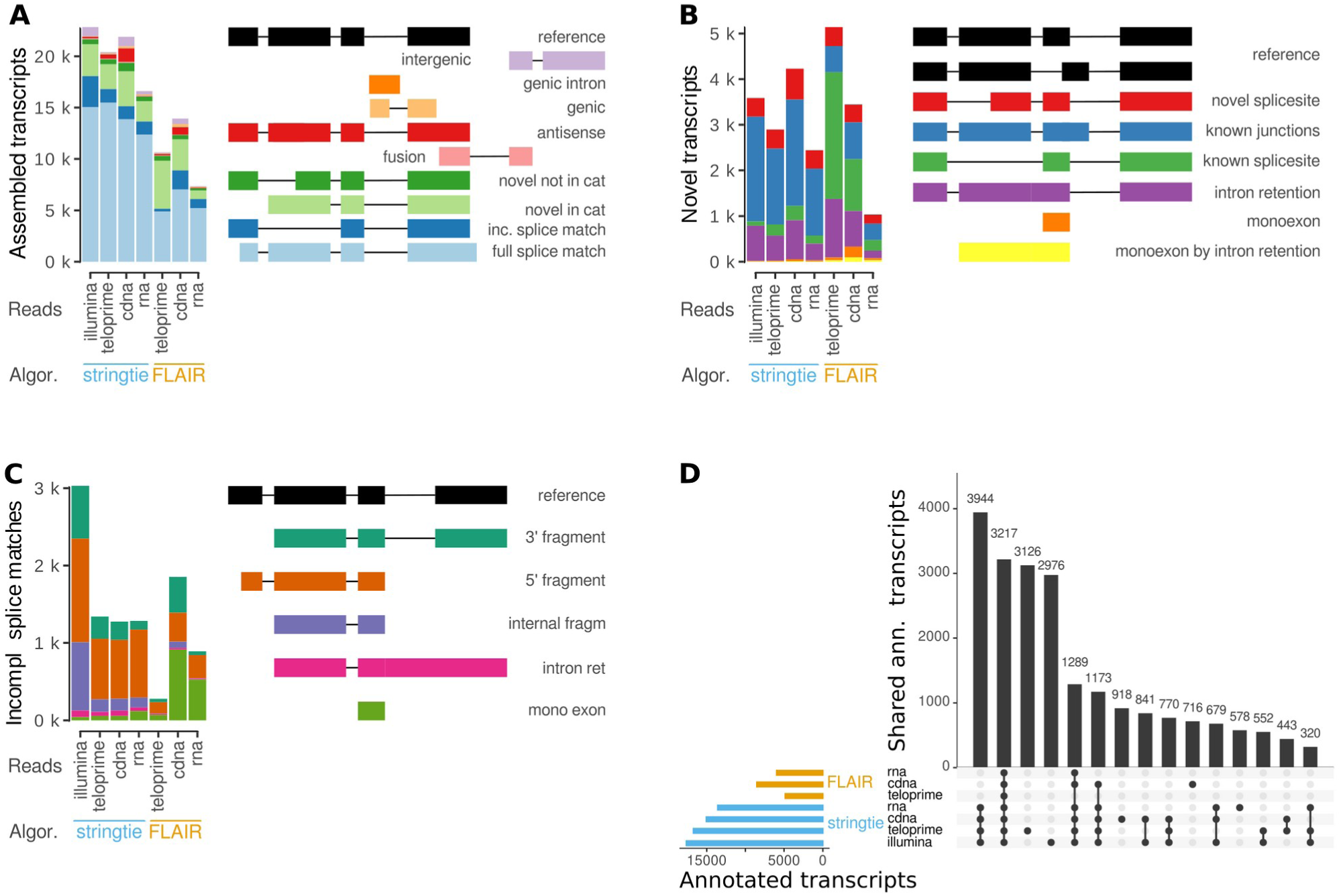
Reannotation analysis of short and long -read sequencing protocols stratified by transcript assembler. **A** Overview of different structural categories. **B;C** Subclassification within the novel transcript and incomplete splice match categories. **D** Overlap between transcripts in the reference annotation (GENCODE M22) correctly reannotated by the different transcriptome assemblers and sequencing methods (includes transcripts falling into the full splice match and incomplete splice match categories as detailed in A).

### Differential transcript usage analysis unravels gene expression alterations upon cold exposure in iBAT

Differential gene expression analysis lacks the sensitivity to detect changes at the transcript-level caused by e.g, alternative transcription start sites (TSS) or alternative splicing (De Paoli-Iseppi et al., 2021). To overcome this limitation, we applied differential transcript usage (DTU) analysis to identify genes using different transcripts in cold-activated compared to inactive iBAT. Reliable identification of DTU depends critically on both the accuracy of the transcript expression quantifications as well as the transcriptome annotation. Therefore, we investigated combinations of transcript quantitation (Illumina or direct cDNA counts) and reannotation algorithms (Fig. 7A). As observed for the differential gene expression analysis (Fig. 5A), a higher number of DTUs were identified when Illumina counts were used to assess transcript quantification (Fig. 7A). We found little overlap between stringtie and FLAIR reannoations in line with other reports (Ringeling et al., 2022). Among the genes with significant isoform switches between cold-activated BAT compared to the controls, we observed phosphodiesterase 4D (*Pde4*d), regulating levels of the signalling intermediate cAMP, which activates lipolysis, glucose uptake, and thermogenesis in brown adipocytes (Reverte-Salisa et al., 2019); the thermogenesis regulating hydrolase androgen dependent TFPI regulating protein (*Adtrp; P. Li et al., 2022)* and regulators of fatty acid metabolism (acyl-CoA synthetase long-chain family member 5; *Acsl5*), glycolysis (Aldolase A; *Aldoa*), protein sorting (endoplasmic reticulum-golgi intermediate compartment 1; *Ergic*1), lipid synthesis (MLX interacting protein-like; *Mlxip*l), beta-oxidation (sterol carrier protein 2, liver; *Scp2*) and protein cysteinylation (cysteinyl-tRNA synthetase gene; *Cars2; Akaike et al., 2017;* Fig. 7C-E and S7). qPCR analysis on iBAT from control and cold-treated mice using primer sets specific for the individual transcripts corroborated our DTU analysis, thus validating the isoform regulation (Fig. 7B). Incorporation of histone modification chromatin immunoprecipitation (ChIP)-Seq peaks into the FLAIR algorithm revealed that *Cars2* is transcribed from two different TSS, i.e. the ‘canonical’ TSS and an alternative gene-internal TSS, giving rise to either the full-length transcript (*Cars2-*FL) or a 5’ truncated transcript (*Cars2-*AT; Fig. S6). Visual inspection of ONT direct cDNA sequencing coverage data indicated that the canonical TSS is dominant at room temperature, while the the alternative TSS is predominantly used in iBAT of cold treated animals (Figure S6A). Cars2 is the mitochondrial cysteinyl-tRNA synthetase, which is important for the translation of mitochondrially encoded genes (Rajendran et al., 2018), but additionally executes a ‘non-canonical’ function in post-translational cysteine and protein persulfidation ultimately affecting mitochondrial respiration (Akaike et al., 2017). Therefore, we analysed the sequences of *Cars2*-FL and *Cars2*-AT to identify whether these alternative transcripts have coding potential using the *Coding-Potential Assessment* (CPA) Tool. CPAT predicted both isoforms to be coding (coding potential > 0.99), annotating the open reading frame correlating with the UniProt reference amino acid sequence to Cars2-FL and predicting a N terminally truncated protein isoform for Cars2-AT using the same open reading frame but missing the first 244 aa (Fig. 7F). As expected for a mitochondrial protein, targetP predicted a mitochondrial localisation signal at the N terminus of the full length CARS2 protein, which was missing in the truncated protein isoform (Fig. 7F). The truncated CARS2 protein isoform was further predicted to lack parts of the conserved binding sites for both Zn^2+^ and pyridoxal phosphate (PLP), indicating a potential lack of catalytic function (Fig. 7F). Using single guide RNAs (sgRNA) targeting either the canonical or the alternative promoter we aimed to specifically overexpress Cars2-FL and Cars2-AT in wt1-SAM immortalised brown adipocytes (Fig. 7G). Using a sgRNA targeting the alternative promoter, it was possible to induce the expression of *Cars2*-AT 10 to 30 times, both with and without β-adrenergic stimulation (p = 0.005 and 0.003) without confounding effects on *Cars2*-FL expression (Fig. 7G). Thus, these results demonstrate that long-read DTU can identify and quantify biologically relevant changes in isoform usage.

**Fig. 7.**
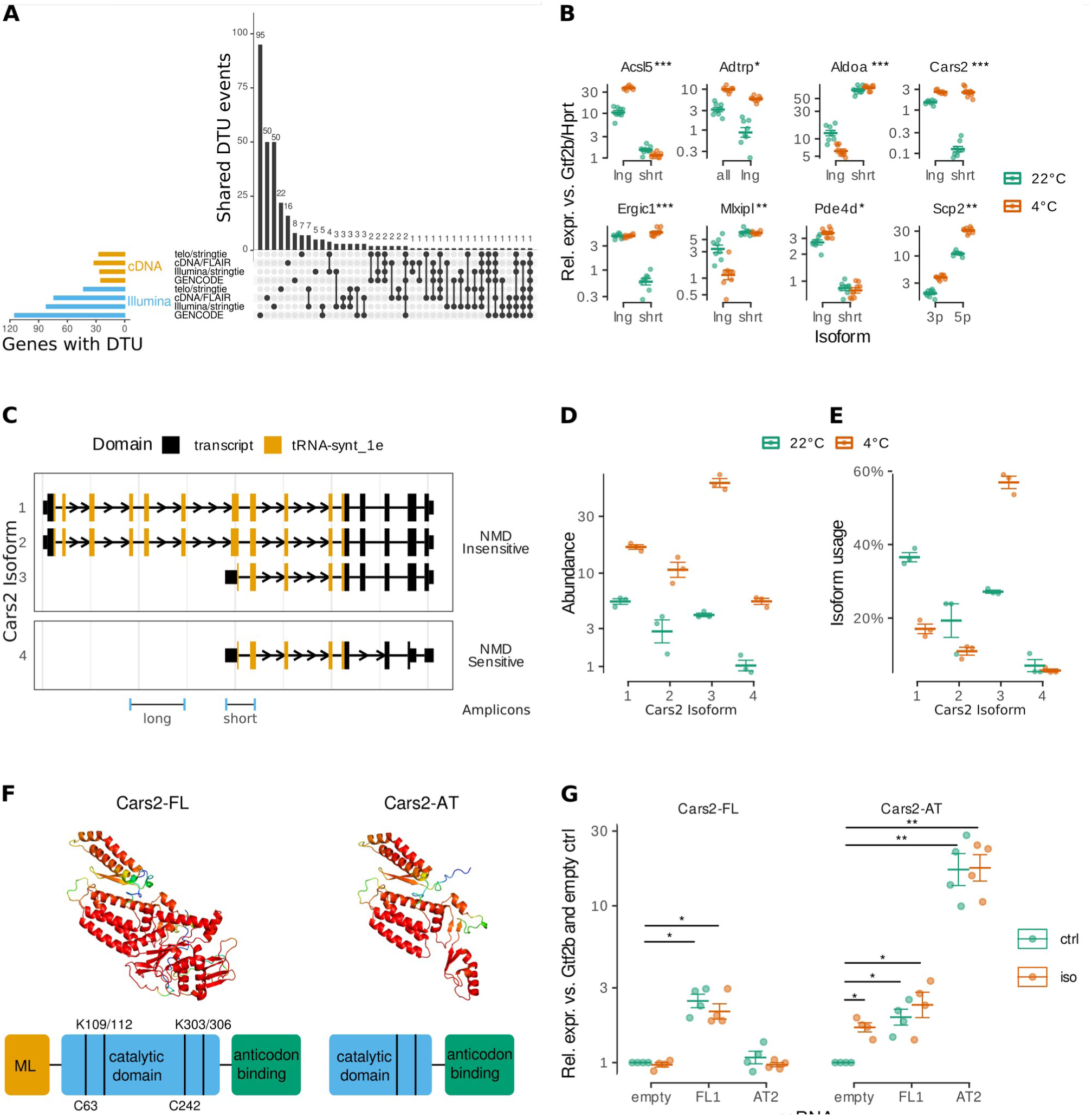
**A** Overlap DTU events detected by different combinations of datasets used for transcript annotation and sequencing. **B** qPCR validation of selected DTU events in murine iBAT from control (22°C) or cold (4°C) treated mice. **C** Structure of expressed *Cars2* isoforms. Arrows show direction of transcription. Narrow lines show intronic regions (not to scale). Exons displayed as boxes. Taller exonic boxes are coding regions, shorter boxes are 5′ and 3′ UTR regions. Colours represent identified protein domains. **D;E** Abundance (normalised counts) and isoform usage of expressed *Cars2* isoforms. **F** Schematic representation of the general structure of Cars2 showing domains and residues important for catalytic activity (modified from (Akaike et al., 2017) and prediction of secondary and tertiary structures of the full length CARS2 and the predicted truncated protein isoform by LocalColabFold. The colour code depicts model confidence; green is low, red is high. **G** Expression of Cars2 isoforms in wt1-SAM brown adipocytes after targeting transcript specifics promoters using sgRNAs.

## 6. Discussion

In this study, we assessed three different ONT long-read sequencing protocols as well as Illumina short-read sequencing for differential gene expression, transcriptome reannotation and differential transcript usage analysis giving us unprecedented views on transcript diversity in murine brown adipose tissue. While studies of differentially expressed genes have provided much of our current understanding of molecular mechanisms controlling BAT function (Seale, 2015; Shapira & Seale, 2019; W. Wang & Seale, 2016), determining genes where functionally distinct alternative transcripts change between BAT activity states is an interesting gene-regulatory mechanism to uncover. Feature quantification using direct RNA and direct cDNA protocols correlated well with Illumina on gene level, and to a lesser extent on transcript level, as described (Sessegolo et al., 2019). Direct RNA and direct cDNA sequencing showed an even higher correlation on gene level which was only slightly reduced when quantifying transcripts, suggesting that *(i)* reverse transcription to cDNA has a limited impact on transcript quantification and *(ii)* long reads give better estimates of transcript abundances, as they more often unambiguously map to a single transcript. In fact, direct RNA as well as direct cDNA even outperform Illumina in terms of accuracy of transcript quantification and differential expression, which has been attributed at least in part to the lack of GC content bias (Chen et al., 2021; Oikonomopoulos et al., 2016, 2020; Sessegolo et al., 2019). The key challenges with direct RNA sequencing are the large amount of input RNA required, higher error rate as compared with cDNA sequencing and the lack of multiplexing options (Chen et al., 2021; Soneson et al., 2019). PCR amplification protocols such as the TeloPrime typically produce higher sequencing depth than PCR-free methods, increasing coverage which is required for accurate identification of alternative transcripts (Chen et al., 2021; Glinos et al., 2022). However, the TeloPrime method overestimated the abundance of highly expressed and underestimated the abundance of lowly expressed features, caused in part by the inherent PCR amplification step compromising transcript diversity (Chen et al., 2021; Sessegolo et al., 2019).

Comparison of the transcriptome reannotation methods showed that when using the same input data, stringtie surpasses FLAIR in terms of the number of correctly reassambled transcripts from the reference annotation (full and incomplete splice matches; Fig. 6A), in line with previous reports (Kovaka et al., 2019). In contrast to stringtie, FLAIR incorporates ChIP/CAGE-Seq data which can be high value because it discriminates between true internal TSS, and artefacts from 5’ degraded RNA, which importantly allowed us to identify a novel 5’ truncated *Cars2* transcript isoform (*Cars2*-AT) highly induced in iBAT of cold-treated mice and predicted to encode a N-terminally truncated protein (Fig. S6). Cars2 has recently been reported to be involved in sulphur metabolism, which is of importance for mitochondrial morphology and BAT function (Akaike et al., 2017; Soriano et al., 2018). The functional significance of the novel *Cars2*-AT reported here, the predicted changes in localization, and its role in thermogenesis remain to be experimentally defined. Our brown adipocyte cell model overexpressing *Cars2*-AT and *Cars2*-FL by CRISPR/Cas9 mediated activation of the respective endogenous promoter (Fig. 7G) will be a valuable tool to answer these questions; and will also allow us to test whether Cars2-AT may have a dominant negative regulatory role in Cars2 expression, as observed for other truncated protein isoforms (Gervois et al., 1999; Tomita et al., 2003). TeloPrime’s strategy for enrichment of full-length transcripts allowed us to identify several novel alternative transcripts produced from the same gene with presumably important functional consequences on protein structure, culminating on thermogenic β3-adrenergic receptor (AR) mediated cAMP signaling (Fig. 7B and C): Cellular cAMP levels are also regulated cAMP-specific phosphodiesterases (PDEs) and *Pde4d* regulates lipolysis and thermogenic gene expression (Vezzosi & Bertherat, 2011). Cold-activated induction of *Pde4d*-long might lead to reduced thermogenic cAMP signalling since *Pde4d*-long controls cAMP levels negatively in a spatial manner due to UCR motifs present in the long *Pde4d* isoform (Lynch et al., 2007). Another factor regulating the thermogenic β3-AR signalling cascade is *Adtrp*; *Adtrp*-deficient mice are cold-intolerant and have defective thermogenesis (P. Li et al., 2022). Thus, cold-activated induction specifically of the longer (enzymatically active) Adtrp isoform shown here might enhance BAT function. Peroxisome derived lipids are required for brown fat-mediated thermogenesis through regulation of cold-induced mitochondrial fission (Kleiboeker & Lodhi, 2022). The cold-induced switch in the peroxisome lipid transfer protein *Scp2* in favour of the longer (and enzymatically active) isoform (Fig. 7B) may thus support a peroxisome-to-mitochondria lipid signalling hub, supporting mitochondrial uncoupling and thermogenesis (N. C. Li et al., 2016). Proper BAT thermogenic function requires cellular protein quality control and removal of misfolded proteins (Bartelt et al., 2018). We here identified DTU in the protein sorting gene *Ergic1 (Joshi et al., 2017)* with the shorter isoform being specifically increased upon cold activation (Fig. 7B;C) which may thus participate in the homeostatic adaptation of BAT to cold stress involving the ER stress response.

The key transcription factor regulating *de novo* lipogenesis *Mlxipl (a.k.a. ChREBP*) inhibits BAT thermogenesis and downregulates expression of genes involved in mitochondrial biogenesis and respiration (Wei et al., 2020). Based on our DTU analysis of *Mlixpl*, we speculate that the observed downregulation of the long, intact *Mlixpl* isoform specifically in cold-activated iBAT alleviates *Mlixpl* inhibitory effects of on BAT thermogenesis. Fatty acid oxidatiocyl-CoA synthetases such as *Acsl5* regulate fatty acid trafficking and metabolism (Bowman et al., 2016). Acsl5 activity increases adiposity, decreases Ucp1 expression and energy expenditure in mice (Bowman et al., 2016). Here, we show DTU in *Acsl5* giving rise to two transcripts differing in their 5’UTR but otherwise identical protein domain structure (Fig. 7B;C) with the longer isoform being the almost exclusively expressed variant in cold, suggesting differential control of translation efficiency of this fatty acid channelling enzyme in response to cold challenge in BAT. A similar scenario may be true for the glyceroneogenic enzyme Aldoa, which controls the cellular levels of glycerol-3-phosphate (G3P) shown to be increased upon cold exposure in BAT of mice (Moura et al., 2005). Thus, a systematic characterization of isoform-level variation and complexity in activated BAT as described here will help understand how isoforms might contribute to the regulation of BAT function.

## 7. Material and methods

### Animal experiments

Unless otherwise stated, mice were kept at 22°C to 24°C on a regular 12 h light cycle with *ad libitum* access to food (Altromin 1324, Altromin Spezialfutter GmbH & Co. KG, Lage, Germany) and water. 20 weeks old male wild type C57BL/6N mice were singly housed at 4 °C for a period of 24 h prior to harvesting adipose tissues.

### RNA isolation

Whole frozen iBAT samples were homogenised in 1 ml TRIsure (Bioline, Memphis, Tennessee, USA) per animal using a tabletop homogeniser (FastPrep-24 5G, MP Biomedicals, Irvine, California, USA). RNA was isolated by phenol chloroform extraction and alcohol precipitation as described by (Workman et al., 2019).

### Illumina RNA sequencing

The NEBNext Ultra II Directional RNA Library Prep Kit (New England BioLabs, Ipswich, Massachusetts, USA) was used to prepare 50 nt paired-end, strand specific libraries following the manufacturer’s protocol and sequenced on a NovaSeq 6000 (Illumina Inc., San Diego, California, USA) for approximately 25 million reads per library.

### ONT library preparation

For all experiments, sequencing on the GridION platform (ONT, Oxford UK) was performed using FLO-MIN106 R9 flowcells (ONT). Libraries prepared according to the TeloPrime and the direct cDNA protocol were sequenced on two different FLO-MIN106 R9 flow cells to examine sequencing variability.

### poly(A) enrichment

RNA used for ONT sequencing was poly(A+) selected in two consecutive rounds using oligo(dT) beads (GenElute mRNA Miniprep Kit, Sigma MRN10, MilliporeSigma, Burlington, Massachusetts, USA) following the manufacturer’s recommendations. Subsequently, RNA was alcohol precipitated using sodium acetate and glycogen following the protocol from the Ribo-Zero rRNA Removal Kit (Illumina).

### TeloPrime libraries

The TeloPrime Full-Length cDNA Amplification Kit (Lexogen, Vienna, Austria) was used to select for full length mRNAs with intact 5’ CAPs from 7 ng poly(A+) RNA. The resulting cDNA was PCR amplified with SYBR Green I (MilliporeSigma), TeloPCR enzyme mix and 3’ and 5’ primers (RP: 5’-TCTCAGGCGTTTTTTTTTTTTTTTTTT-3’ and FP: 5’-TGGATTGATATGTAATACGACTCACTATAG-3’) to determine the optimum cycle numbers for the large-scale PCR to generate enough material for long-read sequencing. The determined cycle number of 27 was applied for large scale PCR in the absence of SYBR Green I followed by processing of 400 ng of the cDNA with the SQK-LSK109 ligation sequencing kit (ONT) and the EXP-NBD104 barcoding kit (ONT) following manufacturer’s instructions.

### Direct cDNA libraries

Libraries were prepared from 100 ng poly(A+) RNA using the SQK-DCS109 direct cDNA sequencing kit (ONT) and the EXP-NBD104 barcoding kit (ONT) according to manufacturer’s protocol.

### Direct RNA libraries

Libraries were prepared from 500 ng poly(A+) RNA using the SQK-RNA002 direct RNA sequencing kit according to manufacturer’s protocol (ONT).

### Reverse transcription and qPCR

RNA was reverse transcribed into cDNA using the High Capacity cDNA Reverse Transcription Kit (Applied Biosystems 4368814, Applied Biosystems, Waltham, Massachusetts, USA) following the manufacturer’s instructions. qPCR was performed in 384 well format in a LightCycler 480 II (Roche, Basel, Switzerland). 4 µl of 1:20 diluted cDNA, 0.5 µl gene specific primer mix (5 µl each) and 4.5 µl FastStart Essential cDNA Green Master (Roche) were amplified using 45 cycles of 25 s at 95 °C, 20 s at 58 °C and 20 s at 72 °C after 300 s at 95 °C initial denaturation. All combinations of primers and samples were run in duplicates and Cq values calculated as the second derivative maximum. Genes of interest were normalised against housekeeper genes using the ΔCq method. The primers used in this study are listed in Tab. S1.

### Long-read alignment and quantification

For the reference-based comparison of ONT library preparation methods, reads were mapped against the transcriptome (GENCODE M22; `-ax map-ont --secondary=no -uf`) and genome (GRCm38.p6; `-ax splice --secondary=no -uf`) using minimap2 (H. Li, 2018). Reads were subsequently filtered for an average PHRED score > 7 using NanoFilt (De Coster et al., 2018). GenomicAlignments (Lawrence et al., 2013) was used to directly extract read level information from BAM files, including transcript level reference sequence, flag based mapping types, mapping position and cigar based aligned length information as described by (Soneson et al., 2019).

### Short-read alignment and quantification

Illumina short-reads were quality filtered using cutadapt (Martin, 2011; `-q 28 -m 30`) and mapped to the genome (GRCm38.p6) using STAR (Dobin et al., 2013). Transcript level quantification was done using salmon in selective alignment mode (GENCODE M22; Patro et al., 2017). Gene level abundance estimation was done using tximport (Soneson et al., 2016).

### Differential gene and transcript expression analysis

Differential gene and transcript expression analysis was done using DESeq2 with H0: log2FC > 0.5 (Love et al., 2014). Adjustment for multiple testing was done using the *s*-value method proposed by Stephens (2017) implemented in apegeglm (Zhu et al., 2018) and *s*-values < 0.05 were considered significant.

### Transcriptome reannotation

Transcriptome reannotation using stringtie (Shumate et al., 2021) was done with a splice junction cutoff (`-j`) of 10 and in presence of the reference annotation (`-G`, GENCODE M22). For ONT long-read runs, the --mix mode was used, additionally providing stringtie with the Illumina-Seq data of the same sample. Afterwards, all reannotations were pooled within the respective library preparation methods using `stringtie --merge` with a coverage cutoff (`-c`) of 3 and only isoforms with a minimum isoform fraction of 5 % per gene were kept (`-f`). No reference annotation was used in the merge step.

Aligned ONT long-reads were corrected by FLAIR correct (Tang et al., 2020) using the splice junction data from STAR from the Illumina-Seq runs of the same samples, filtered for a minimum splice junction coverage of 10. Subsequently, the corrected reads were collapsed into a transcriptome reannotation by FLAIR collapse using a minimum coverage of 3 and setting the `--stringent` flag. In order to mark true transcriptional start sites, a combined bed file from FANTOM5 CAGE peaks and iBAT H3K4me3 peaks was provided (Abugessaisa et al., 2017; Engelhard et al., 2022). For the TeloPrime data, the `-- trust_ends` flag was additionally set.

The reannotated transcripts were compared to the reference annotation using SQANTI2 (Tardaguila et al., 2018)and the overlap between the different sequencing methods was calculated based on the associated transcripts for transcripts either fully or partially matching a reference transcript as described elsewhere in detail (Soneson et al., 2019).

### Analysis of differential transcript usage

Only genes with at least 10 counts in all samples were included in the analysis. Transcripts were filtered for a minimum of 5 counts and 10 % of the counts of the parent gene in half of the samples. Differential transcript usage analysis was done using DRIMSeq and FDR calculated using stageR (Love et al., 2018; Nowicka & Robinson, 2016; Van den Berge et al., 2017). FDR < 0.1 and transcript usage changes > 5 % were considered significant. In order to detect differential transcript usage from qPCR data, a linear model was fitted with isoform and temperature as variables and the interaction term was tested. *p*-values were adjusted for multiple testing using Holm’s method.

### Annotation of transcripts

Annotation of transcript isoforms including open reading frames, nonsense mediated decay (spliceR, Vitting-Seerup et al., 2014) and functional protein domains (pfam, Finn et al., 2016) was done using IsoformSwitchAnalyzeR (Vitting-Seerup & Sandelin, 2019).

Cars2 protein fold predictions were generated in LocalColabFold using standard parameters (Mirdita et al., 2022). Computation of the models was performed on the UCloud interactive HPC system, which is managed by the eScience Center at the University of Southern Denmark.

### Cars2 overexpression in cell culture

wt1-SAM brown preadipocytes were grown in high glucose DMEM supplemented with 10 % fetal bovin serum (FBS) and 1 % penicillin-streptomycin. After reaching confluency, differentiation was induced by 0.5 μM rosiglitazone, 1 nM T3, 1 μM Dexamethasone, 850 nM insulin, 125 μM indomethacine and 500 μM IBMX. Two days later, medium was exchanged for medium supplemented with 0.5 μM rosiglitazone and 850 nM insulin. Afterwards, medium was changed for medium containing 0.5 μM rosiglitazone every second day until reaching full differentiation 7 days after induction.

For *in vitro* gain of function studies using the wt1-SAM cell line, single guide RNAs (sgRNAs) were designed using CRISPick (Doench et al., 2014) and cloned into the sgRNA(MS2) cloning backbone (addgene 61424) as described by (Konermann et al., 2015). To transfect mature adipocytes, 3 μl TransIT and 250 ng plasmid DNA or 1.4 μl (10 μM) in 100 μl Opti-MEM I were pipetted into a well of a 24 well plate. After 15 min, 500,000 cells resuspended in 500 μl Opti-MEM I were added. 24 h later, medium was changed for regular differentiation medium. Guide sequences used were ATTTAGGCATTTGGGCACGG for Cars2-FL and GTGGCTGAACAGATCTGGCC for Cars2-AT.

## 8. Data and software availability

The RNA-Seq data presented in this study has been deposited to GEO and will be made public once this study has been accepted for publication. Reviewers can get access via a reviewer token upon request. Source code for the reproduction of all analyses and plots in this study are available at https://gitlab.com/kiefer.ch/nanoporeibat_new.

## 9. Acknowledgements

JWK and CHE were funded by the University of Southern Denmark and the Danish Diabetes Academy, which is funded by the Novo Nordisk Foundation. ONT sequencing was performed at the Core Facilities of the Medical University of Vienna, a member of VLSI. We thank Markus Jeitler (Core Facilities, Medical University of Vienna) for help with ONT library preparation and sequencing, and Bjørk Ditlev Marcher Larsen and Ajeetha Josephrajan (Department for Biochemistry and Molecular Biology, University of Southern Denmark) for generating and visualising the Cars2 structure predictions, respectively. The wt1-SAM brown preadipocyte cell line as well as the empty sgRNA vector have been generated as described in (Lundh et al., 2017) and were kind gifts of Brice Emanuelli.

## 10. Conflict of interest

The authors declare no competing interest.

## 11. Supplementary Figures

**Fig S1:**
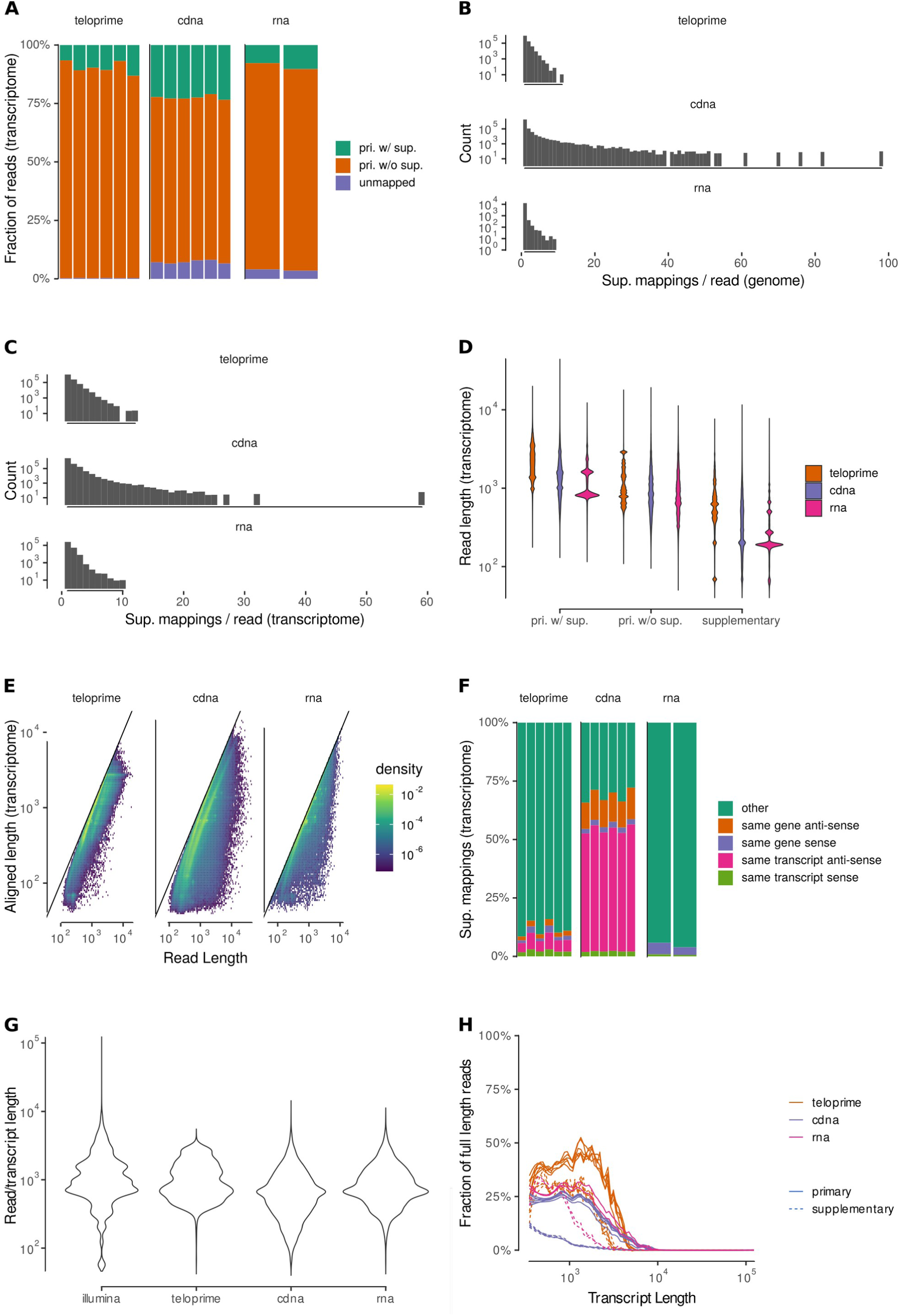
Characterisation of read alignments. **A** Fraction of reads classified by whether the primary alignment against the transcriptome has at least one supplementary alignment attached to it. **B;C** Distribution of supplementary alignments per primary read for genome (B) and transcriptome (C) mapping. **D** Read length distribution for reads aligned to the transcriptome. **E** Aligned length vs read length for primary alignments to the transcriptome. **F** Fraction of supplementary alignments against the transcriptome stratified by their relation to the corresponding primary alignment. **G** Read length distribution of primary alignments to the transcriptome compared to the hypothetical transcript length distribution as estimated from the short-read transcript abundances. **H** Smoothed fraction of full length alignments (> 90 %) mapped to the transcriptome for transcripts > 350 nt.

**Fig S2:**
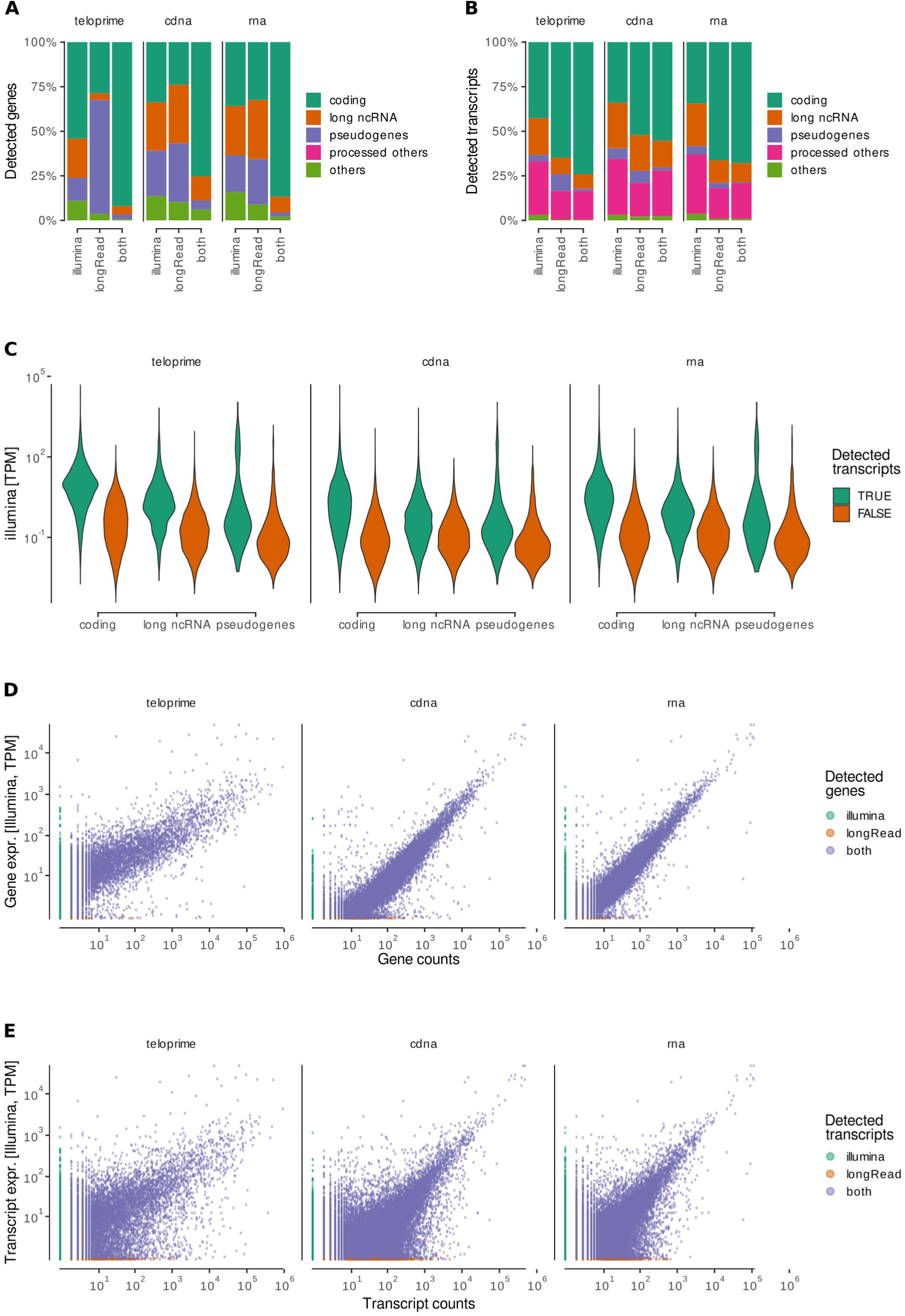
Feature detection. **A;B** Biotypes of genes (A) and transcripts (B) detected by the indicated method. A feature is counted as detected if there is at least one primary alignment to it. **C** Illumina-Seq based abundance of transcripts stratified by whether they are detected by the different ONT library preparation methods or not. **D;E** Expression levels of genes (D) and transcripts (E) stratified by whether they are detected by Illumina, the respective ONT library preparation method, or both.

**Fig S3:**
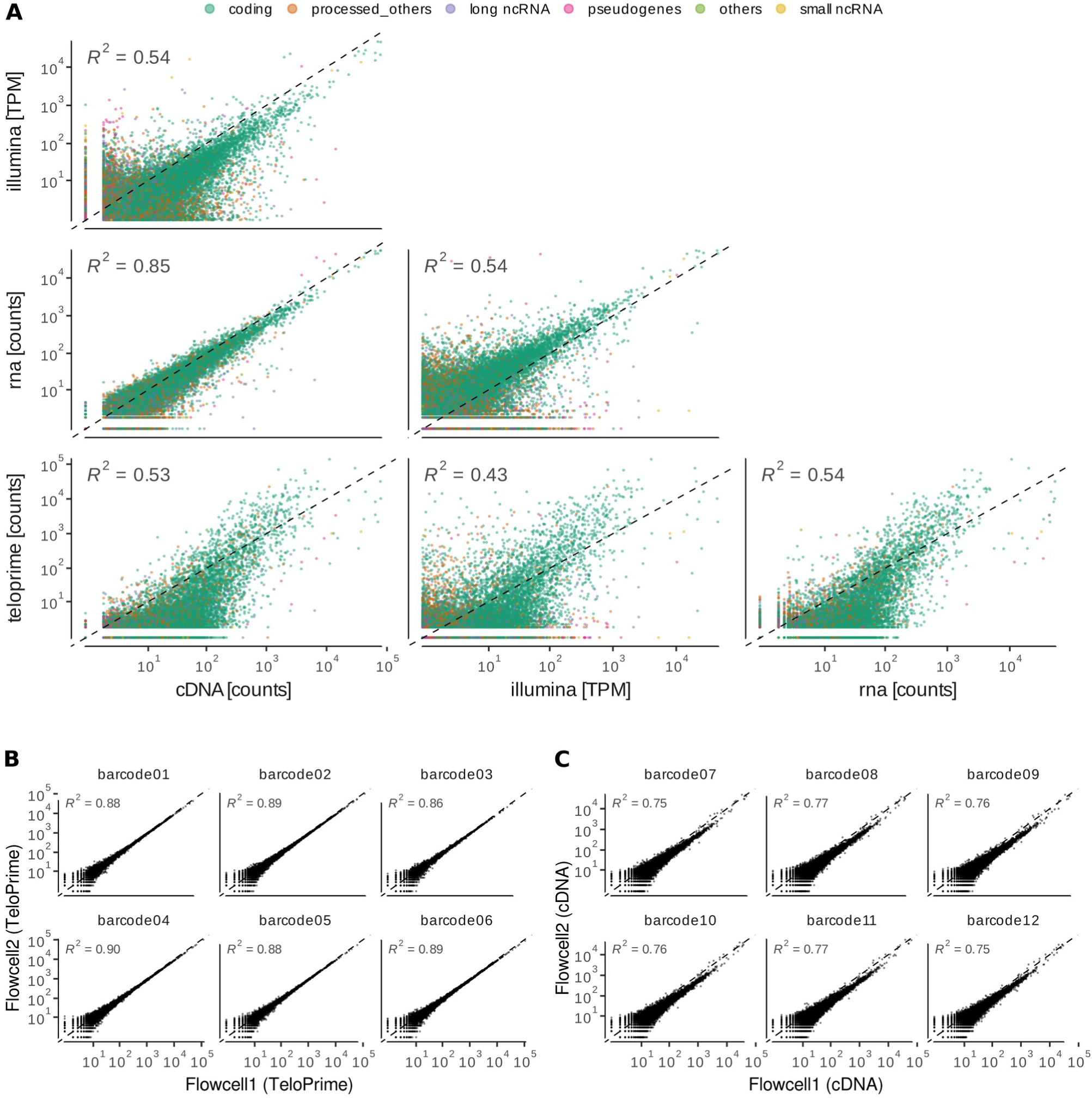
Transcript quantification. **A** Scatter plots showing the correlation in transcript quantification between the different sequencing methods. **B;C** Sample wise correlation analysis of transcript quantification between two different flow cells for the TeloPrime (B) and direct cDNA (C) methods.

**Fig S4:**
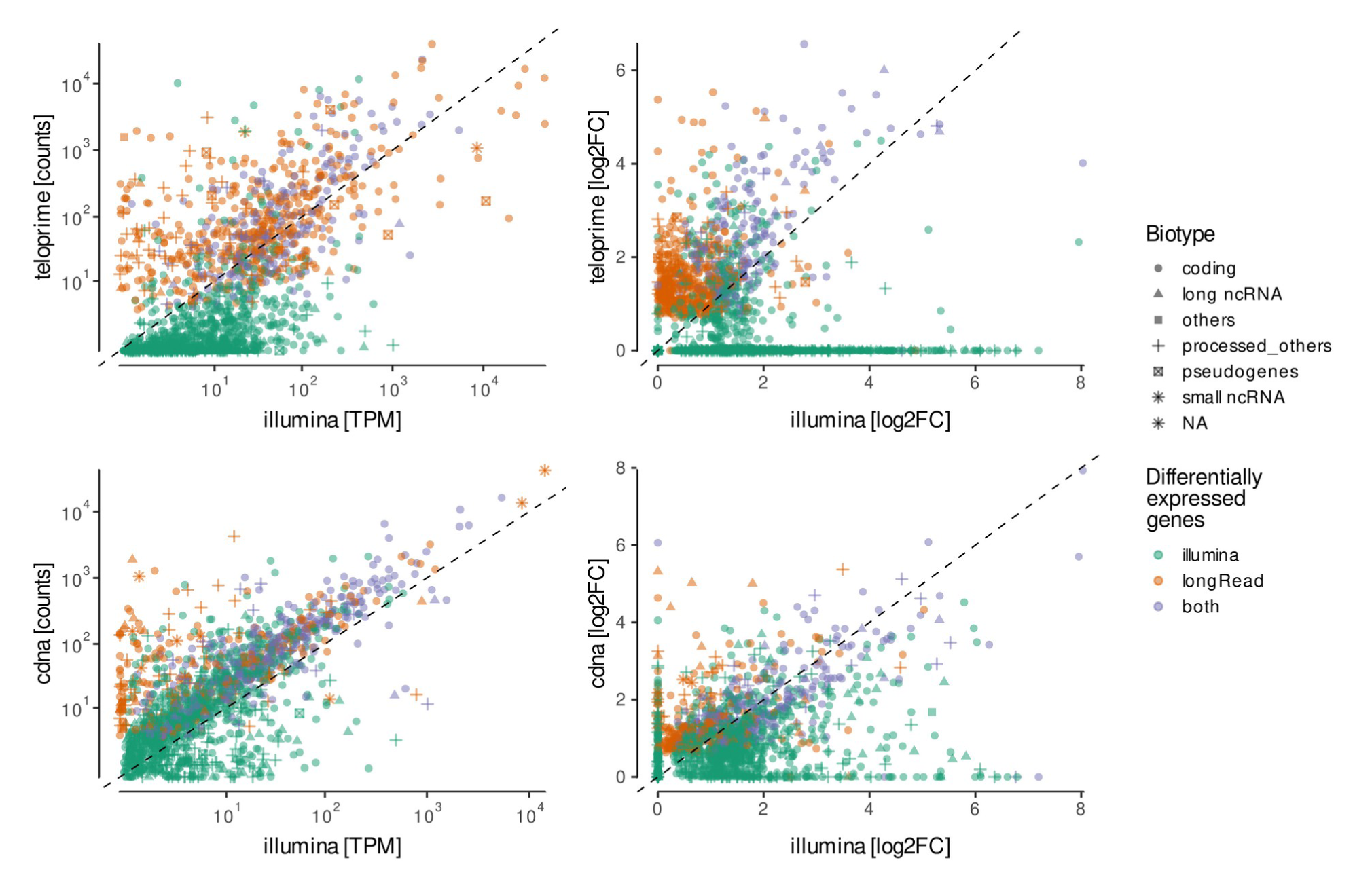
Differential transcript expression analysis. Comparison of expression levels and fold changes of transcripts showing significant differential expression between direct cDNA/TeloPrime and Illumina sequencing.

**Fig S5:**
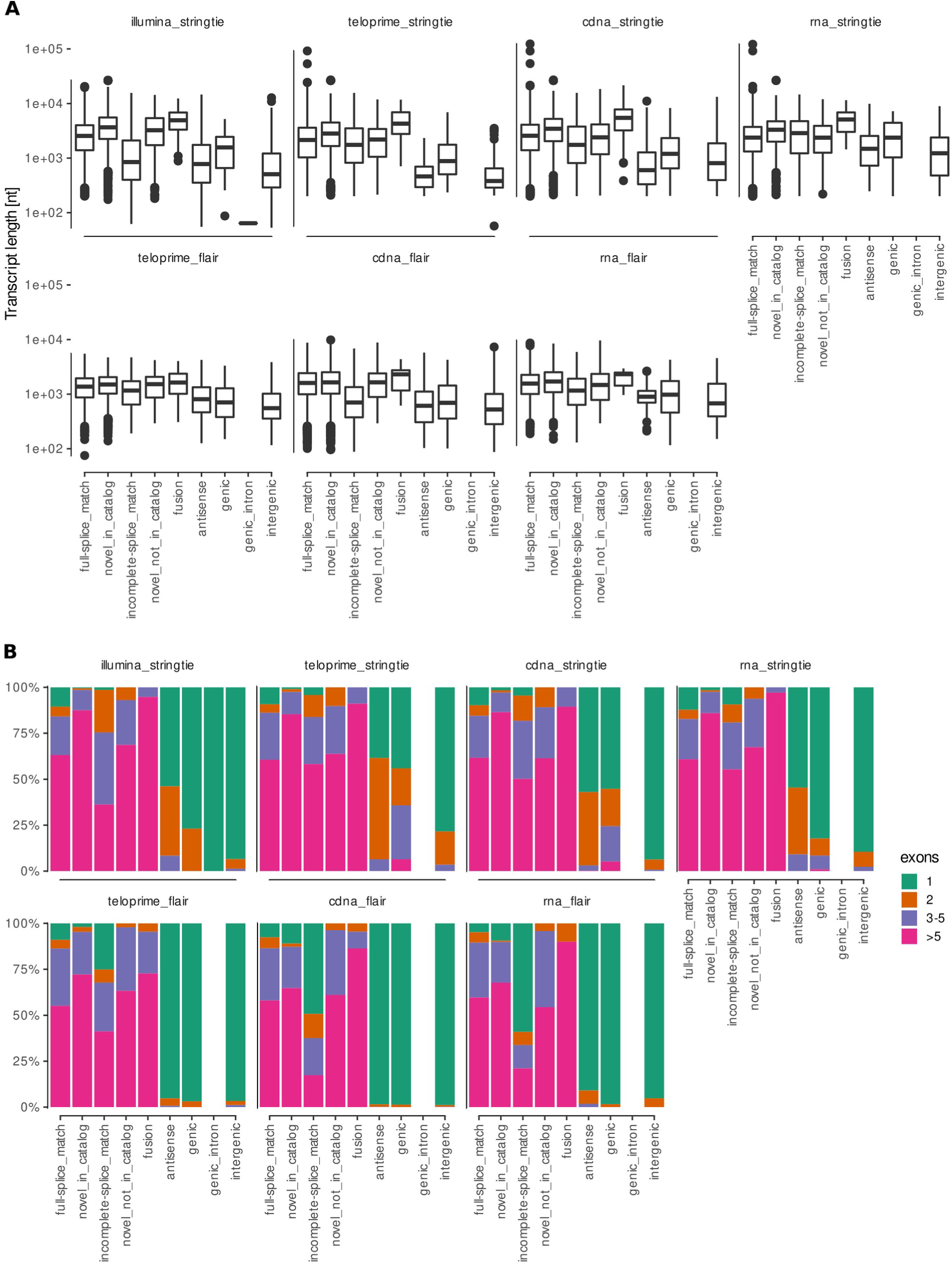
Reannotation Analysis. Transcript length distributions (A) and distributions of number of exons per reannotated transcript (B) compared for different structural categories and different reannotation methods. Center lines represent the median; hinges represent first and third quartiles; whiskers the most extreme values within 1.5 interquartile range from the box.

**Fig S6:**
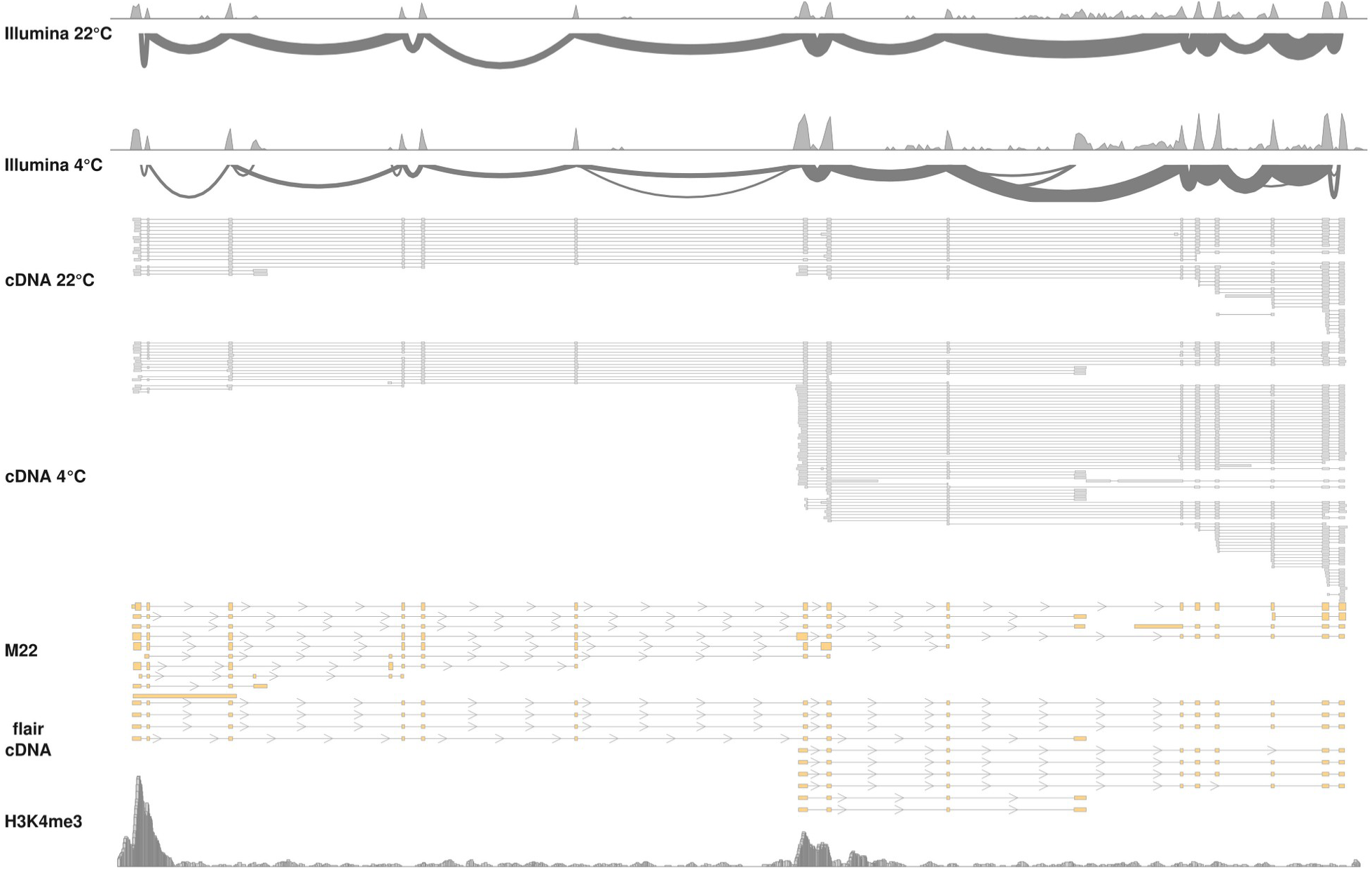
Detailed analysis of the Cars2 locus. Browser tracks of transcripts annotated for the *Cars2* locus in murine iBAT. Shown are the coverage (log10(x+1)) and splice junction usage from two representative Illumina sequencing runs, long-read alignments from two representative direct cDNA sequencing runs, as well as a the reannotated transcripts from FLAIR compared to the reference annotation (GENCODE M22) and histone ChIP-Seq peaks marking the TSSs. Note the diversity in Cars2 transcripts and their usage between treatments.

**Fig S7:**
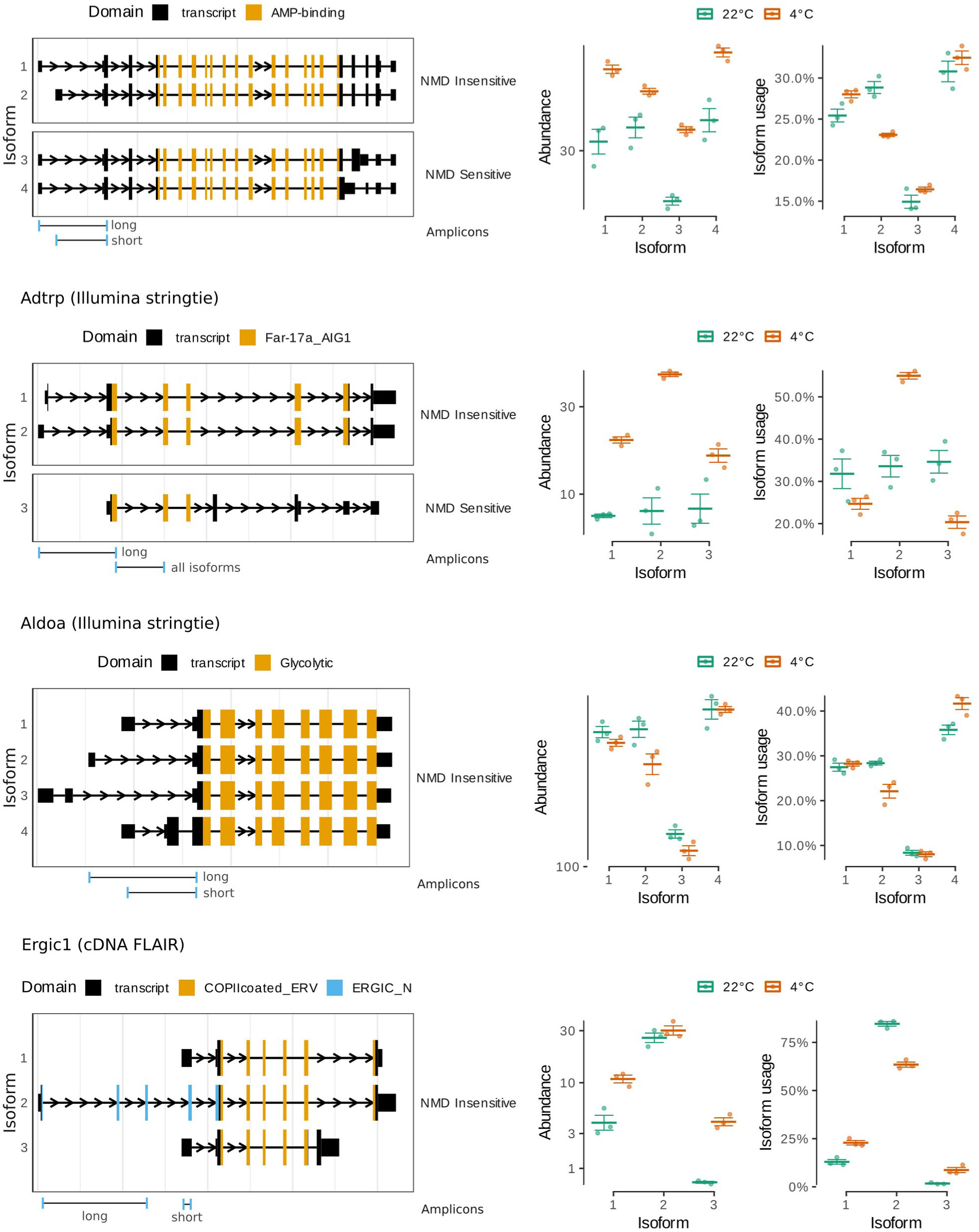
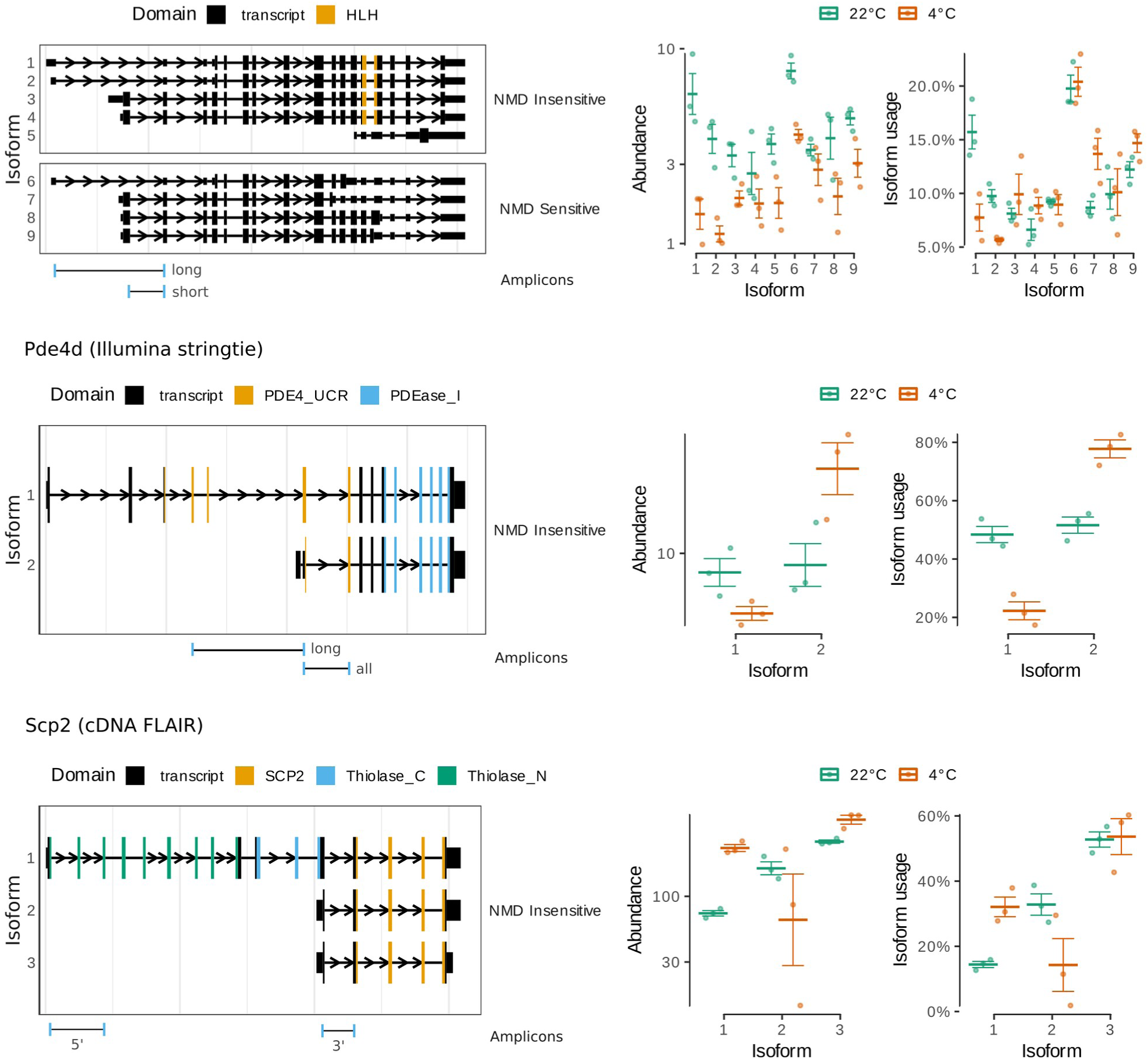
Differential transcript usage in cold-activated murine iBAT. Structure of expressed isoforms of genes showing differential isoform usage detected in the indicate dataset.

## 12. Supplementary tables

**Tab. S1.**
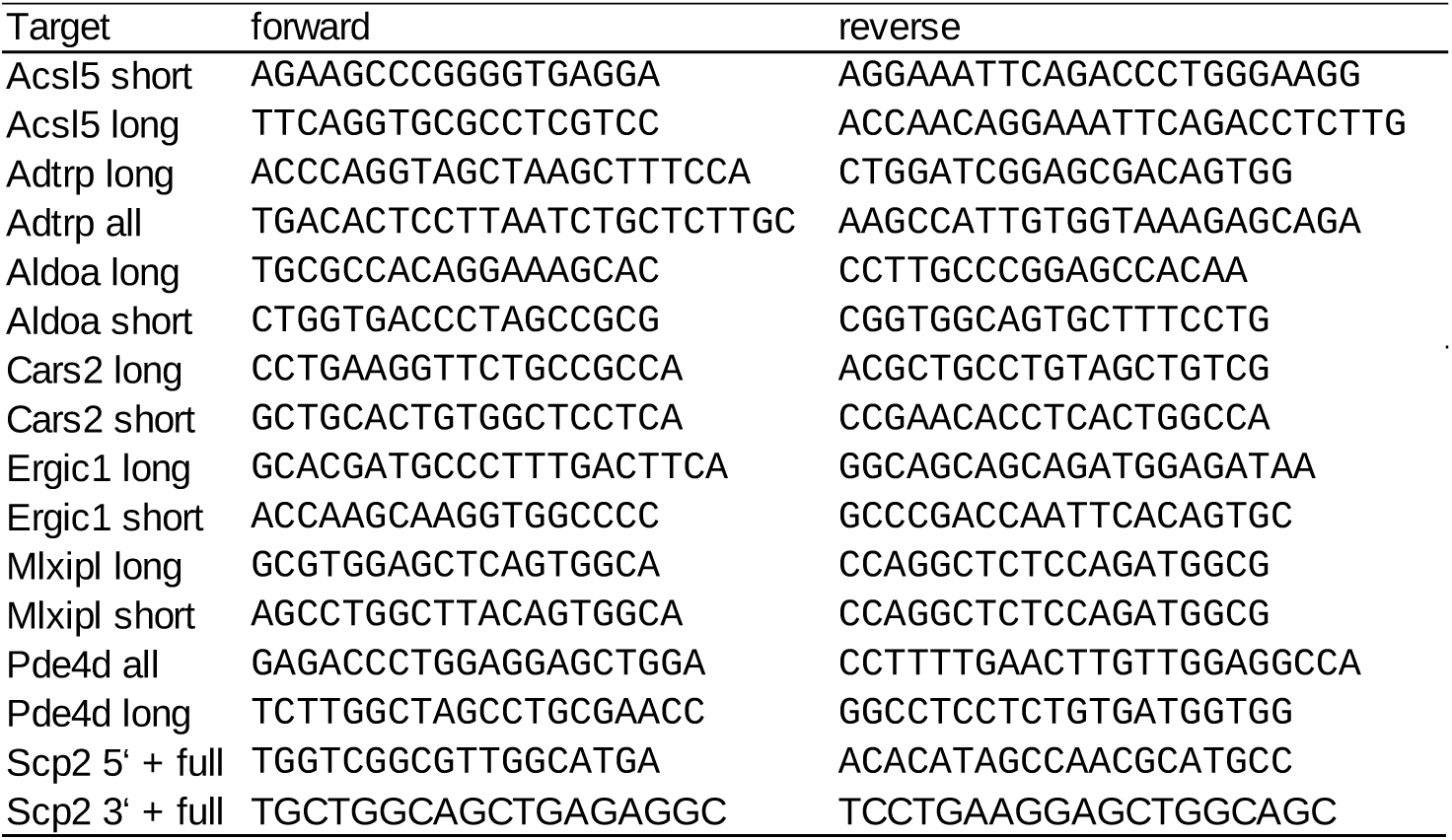
qPCR primers.

## Abbreviations

BAT: brown adipose tissue
cDNA: complementary DNA
DTU: differential transcript usage
iBAT: interscapular brown adipose tissue
lncRNA: long non-coding RNA
mRNA: messenger RNA
ONT: Oxford Nanopore Technologies
PCR: polymerase chain reaction
qPCR: quantitative polymerase chain reaction
RNA: ribonucleic acid
TSS: transcription start site

## Notes

### Competing Interest Statement

The authors have declared no competing interest.

